# Divergence in dimerization and activity of primate APOBEC3C

**DOI:** 10.1101/2021.07.13.452235

**Authors:** Amit Gaba, Mark A. Hix, Sana Suhail, Ben Flath, Brock Boysan, Danielle R. Williams, Tomas Pelletier, Michael Emerman, Faruck Morcos, G. Andrés Cisneros, Linda Chelico

## Abstract

The APOBEC3 (A3) family of single-stranded DNA cytidine deaminases are host restriction factors that inhibit lentiviruses, such as HIV-1, in the absence of the Vif protein that causes their degradation. Deamination of cytidine in HIV-1 (-)DNA forms uracil that causes inactivating mutations when uracil is used as a template for (+)DNA synthesis. For APOBEC3C (A3C), the chimpanzee and gorilla orthologues are more active than human A3C, and the Old World Monkey A3C from rhesus macaque (rh) is not active against HIV-1. Multiple integrated analyses determined why rhA3C was not active against HIV-1 and how to increase this activity. Biochemical, virological, and coevolutionary analyses combined with molecular dynamics simulations showed that the key amino acids needed to promote rhA3C antiviral activity also promoted dimerization. Although rhA3C shares a similar dimer interface with hominid A3C, the key amino acid contacts were different. Overall, our results determine the basis for why rhA3C is less active than human A3C, establish the amino acid network for dimerization and increased activity, and track the loss and gain of A3C antiviral activity in primates. The coevolutionary analysis of the A3C dimerization interface provides a basis from which to analyze dimerization interfaces of other A3 family members.

## Introduction

Host restriction factors act as a cross-species transmission barrier for the Simian and Human Immunodeficiency viruses, SIV and HIV (Gaba, Flath, & Chelico, 2021). Overcoming these barriers by evolution of virally-encoded ‘accessory proteins’ which suppress these restriction factors has characterized successful adaptation of the primate lentiviruses to new hosts (Harris, Hultquist, & Evans, 2012). In some cases, such as the infection of chimpanzees with SIV from Old World Monkeys (OWM) these adaptions to a new host has been accompanied by major changes in the viral genome, which also facilitated the transmission to humans as HIV-1 (Etienne, Hahn, Sharp, Matsen, & Emerman, 2013). One of the major barriers to cross-species transmission of SIVs to a new host is the family of APOBEC3 (A3) host restriction factors that in primates constitutes a minimum of seven enzymes, named A3A through A3H, excluding E (Uriu, Kosugi, Ito, & Sato, 2021). The A3 family are single-stranded (ss) DNA cytidine deaminases that can inhibit retroelements, such as LINE-1, retroviruses, such as HIV and SIV, and other viruses (Adolph, Love, & Chelico, 2018; Arias, Koyama, Kinomoto, & Tokunaga, 2012; Cheng et al., 2021; Delviks-Frankenberry, Desimmie, & Pathak, 2020; Xu, Byun, & Dudley, 2020). They belong to a larger family of APOBEC enzymes that have diverse roles in immunity and metabolism (Salter, Bennett, & Smith, 2016).

For restricting HIV-1 replication, A3 enzymes first need to become packaged in the budding virion (Adolph et al., 2018; Xu et al., 2020). When these newly formed virions infect the next target cell, the packaged A3 enzymes deaminate cytidines on single-stranded regions of the (-) DNA to uridines during reverse transcription (Adolph et al., 2018; Xu et al., 2020). Uracil is promutagenic in DNA since it templates the addition of thymine and results in G to A hypermutations on the (+)DNA when it is synthesized using a deaminated (-)DNA as a template (Adolph et al., 2018; Xu et al., 2020). This resulting double-stranded (ds) DNA provirus can become integrated but is non-functional (Adolph et al., 2018; Xu et al., 2020). Some A3 enzymes can also physically inhibit HIV reverse transcriptase activity (Iwatani et al., 2007; Pollpeter et al., 2018). To counteract the actions of A3 enzymes, HIV encodes a protein Vif that becomes the substrate receptor of an E3 ubiquitin ligase with host proteins CBF-β, Cul5, EloB, EloC and Rbx2 to cause polyubiquitination and degradation of A3 enzymes via the 26S proteasome (Hu, Knecht, Shen, & Xiong, 2020). The members of human A3 family exhibit considerable variation in their antiviral activity. A3G can restrict HIV-1 ΔVif replication the most, with A3H, A3F, and A3D having decreasing abilities to restrict HIV-1 ΔVif (Adolph, Ara, et al., 2017; Ara, Love, & Chelico, 2014; Chaipan, Smith, Hu, & Pathak, 2013; OhAinle, Kerns, Li, Malik, & Emerman, 2008). A3A and A3B can potently restrict endogenous retroelements and DNA viruses, respectively, but not HIV-1 (Arias et al., 2012; Cheng et al., 2019; Richardson, Narvaiza, Planegger, Weitzman, & Moran, 2014).

A3C is the least active member of the human A3 family when it comes to restricting HIV-1, HIV-2, and endogenous retroelements (Hultquist et al., 2011; Muckenfuss et al., 2006; Nchioua et al., 2021). However, a polymorphism in A3C that exists in about 10% of African individuals causes a change of Serine 188 to Isoleucine 188, and this increases its HIV ΔVif restriction ability 5- to 10- fold, but it does not reach the level of restriction that A3G can achieve (Anderson et al., 2018; Wittkopp, Adolph, Wu, Chelico, & Emerman, 2016). The increased restrictive activity of A3C S188I is due to acquiring the ability to form a homodimer, which enables processive deamination of HIV (-)DNA (Adolph, Ara, et al., 2017). However, for the common form of human (h)A3C there is also a contribution of a specific loop near the active site of A3C (loop 1) to catalytic activity (Jaguva Vasudevan et al., 2020). The common hA3C is catalytically active, but is unable to significantly deaminate (-)HIV DNA due to a lack of processivity (Adolph, Ara, et al., 2017). Since A3 enzymes only deaminate ssDNA and the availability of ssDNA during reverse transcription is transient, the enzymes must have an efficient way to search for cytidines in the correct deamination motif, e.g., 5’TTC for A3C, in order to maximize deaminations in the short time that the ssDNA is available (Adolph et al., 2018). Interestingly, chimpanzee (c)A3C and gorilla (g)A3C despite having a S188 have equivalent HIV restriction activity to hA3C S188I (Adolph, Ara, et al., 2017). This restriction activity of cA3C and gA3C is also attributed to its ability to form dimers, but through a different amino acid at position 115 (Adolph, Ara, et al., 2017).

In contrast to hominid A3C, the rhesus macaque (rh)A3C has been found to not be able to restrict replication of HIV-1, HIV-2, SIVmac (rhesus macaque), or SIVagm (african green monkey) in the majority of studies (Hultquist et al., 2011; Virgen & Hatziioannou, 2007; Zhang et al., 2016). Paradoxically, the rhA3C contains the I188 that enables activity of hA3C and enhances activity of cA3C, and gA3C (Adolph, Ara, et al., 2017; Wittkopp et al., 2016). Using molecular dynamics (MD) simulations, we found that rhA3C had several amino acid differences that when present in a hA3C backbone disrupted dimerization. Through direct coupling analysis (DCA)/coevolutionary analysis and subsequent MD, virological, and biochemical experiments, we uncovered the series of amino acid replacements required to promote dimerization in rhA3C and therefore promote HIV-1 restriction ability. Analysis of several OWM A3C sequences showed that they did not contain the “right” combination of amino acids for activity against HIV-1 suggesting that the evolutionary pressures that formed OWM A3C were different from hominid A3Cs that are active against lentiviruses. Overall, we form a model for the determinants of A3C antiviral activity and estimate its loss and gain throughout primate evolution.

## Results

### rhA3C is a monomer

Dimerization of hA3C, cA3C, and gA3C has previously been shown to correlate with HIV-1 restriction efficiency (Adolph, Ara, et al., 2017). Since rhA3C has been reported as not being active against HIV (Hultquist et al., 2011; Virgen & Hatziioannou, 2007; Zhang et al., 2016), we hypothesized that this could be due to a lack of dimerization. To determine the overall amino acid similarity, particularly at the dimer interface residues the amino acid sequences for hA3C, cA3C and rhA3C were aligned (Figure 1). The percent amino acid identity for hA3C and cA3C is 98%, but for both hA3C and cA3C compared to rhA3C, it is only 85% and 84%, respectively. The rhA3C does contain an I188, which stabilizes dimerization in hA3C and cA3C (Figure 1) (Adolph, Ara, et al., 2017). However, the other determinant identified for cA3C that promotes activity against HIV-1 is K115. The rhA3C has a different amino acid at this position, M115, which is also different from hA3C that has an N115 (Figure 1). Furthermore, amino acid R44 (and possibly R45) was predicted to interact with K115 in cA3C (Adolph, Ara, et al., 2017), but in rhA3C these amino acids are Q44 and H45.

**Figure 1.**
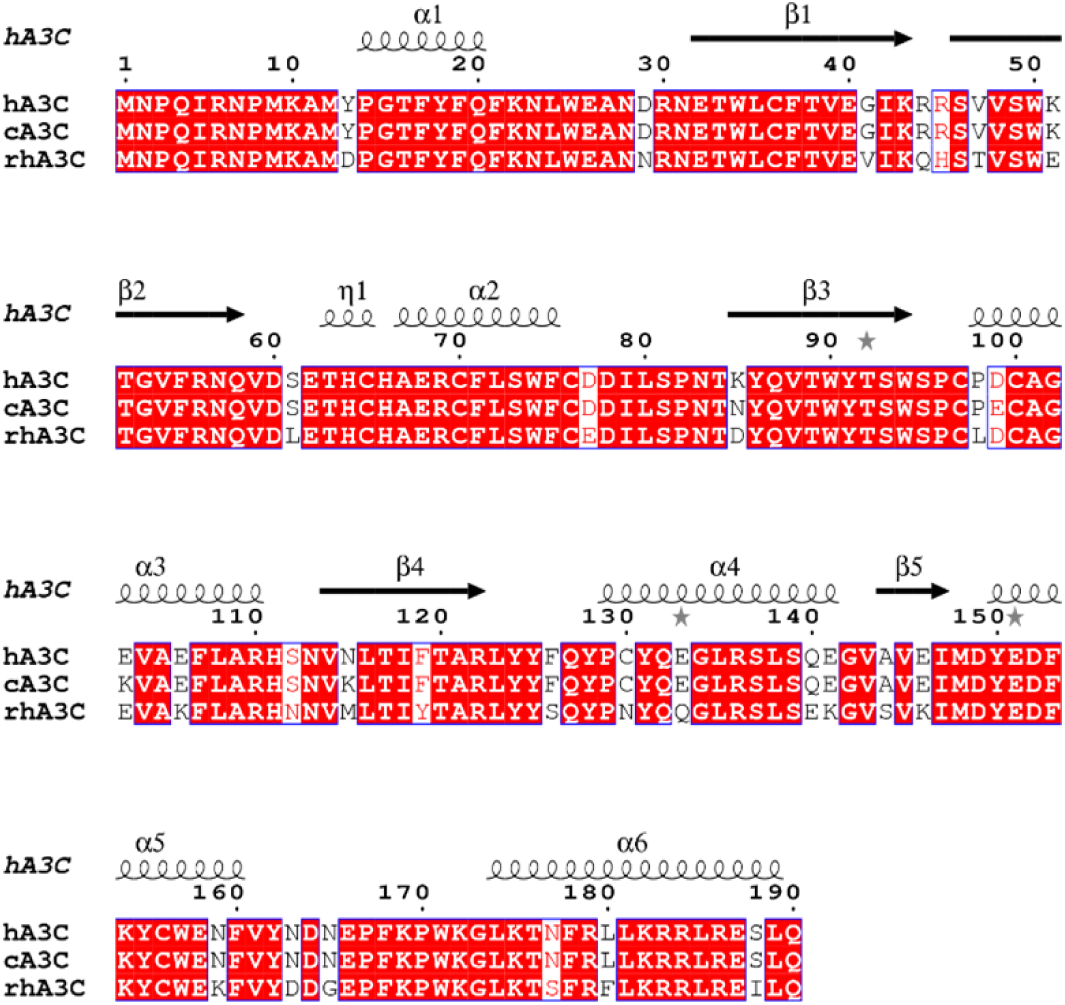
Amino acid differences of rhA3C in comparison to hA3C and cA3C. Sequence alignment and structural analysis of A3C. (**A**) Sequence alignment of hA3C, cA3C and rhA3C with amino acid differences shown in white. The sequence alignment was performed by a Clustal Omega multiple sequence alignment (Sievers et al., 2011) and plotted using the program ESPript (Robert & Gouet, 2014). The α-helices and β-strands are shown above the alignment and are based on the hA3C 3VOW structure (Kitamura et al., 2012).

To determine if these amino acid differences had an effect on rhA3C dimerization, we purified the rhA3C wild type (WT). The rhA3C WT was produced from *Sf*9 cells and we used size exclusion chromatography (SEC) to determine the oligomerization state. We found that the rhA3C WT had only one peak in the chromatogram that was a monomer (M, Figure 2A-B), suggesting that the amino acid differences identified prevent dimerization. We then created rhA3C mutants to make it more hominid-like. We purified rhA3C mutants Q44R/H45R (hA3C and cA3C -like), Q44R/H45R/M115K (cA3C-like), and Q44R/H45R/M115N (hA3C-like). The rhA3C Q44R/H45R and Q44R/H45R/M115K mutants also only had one peak in the chromatogram that was a monomer (M, Figure 2A-B). However, the rhA3C Q44R/H45R/M115N had two peaks demonstrating formation of a dimer (D), but the monomer peak was more predominant (Figure 2A-B).

**Figure 2.**
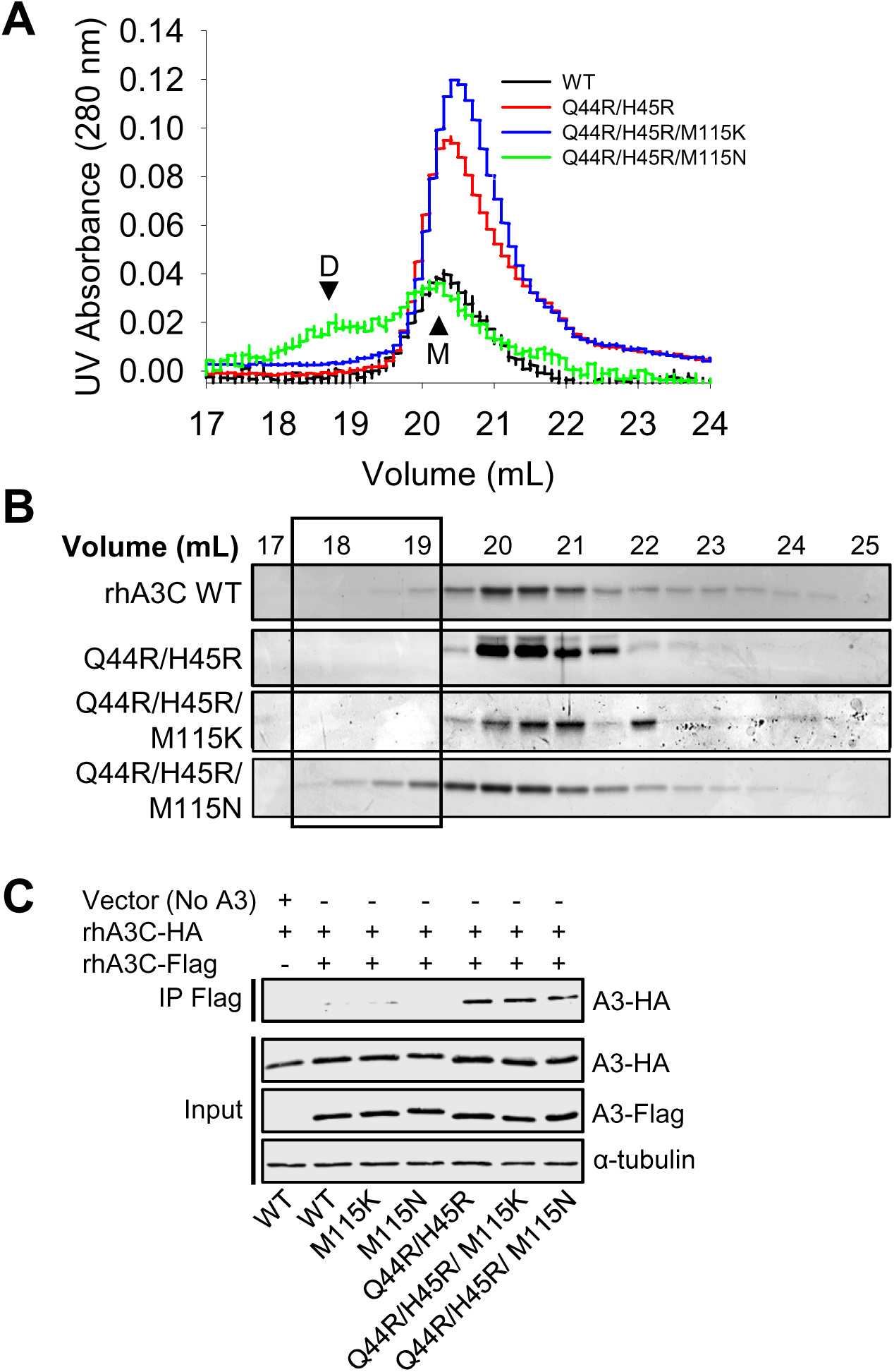
Mutation of rhA3C amino acids to hA3C amino acids enables dimerization. The oligomerization of rhA3C was studied **(A-B)** *in vitro* and **(C)** in cell lysates. **(A-B)** SEC profile for rhA3C WT and rhA3C mutants Q44R/H45R, Q44R/H45R/M115K, and Q44R/H45R/M115N. Elution profiles for all enzymes was composed of a monomer peak (M, ∼20 ml elution volume). Only for the rhA3C Q44R/H45R/M115N there was a larger dimer peak (D, ∼19 mL elution volume). The elution profiles are shown as the **(A)** UV absorbance during SEC elution and **(B)** coomassie stained protein fraction resolved by SDS-PAGE. Box shows where the dimeric fractions eluted. During the determination of the apparent molecular weights in comparison to a series of standards, we found that the A3C retention time in the SEC column did not reflect the actual molecular weight (Figure S1). However, comparison to cA3C confirmed our assignment of monomer and dimer forms and use of Multi-angle light scattering to empirically determine the molecular mass of rhA3C Q44R/H45R/M115K confirmed that all A3C forms were monomeric except Q44R/H45R/M115N (Figure S1). **(C)** Co-immunoprecipitation of rhA3C-3xHA with rhA3C-Flag. The A3C-3xHA and A3C-Flag were transfected in combination and immunoprecipitation of the cell lysates used magnetic anti-Flag beads. Immunoblotting was conducted with antibodies against α-tubulin, HA and Flag. Cell lysate (input) shows the expression of α-tubulin, HA and Flag.

To determine if the rhA3C Q44R/H45R/M115N could also form dimers in cells we used co-immunoprecipitation (co-IP). Due to the propensity of A3s to bind RNA in cells and cell lysates (Smith, 2016), the co-IP was carried out in the presence of RNaseA to ensure that interactions were due to protein-protein contacts and not mediated by an RNA bridge. Using two rhA3C constructs with either a 3xFlag- or 3xHA- tag we carried out a co-IP and found that the rhA3C Q44R/H45R/M115N was able to self-associate in cells. As a control, we also tested the single mutant M115N and this could not immunoprecipitate, suggesting that the Q44R/H45R change enabled dimer formation (Figure 2C). Consistent with this, the rhA3C self-interaction was also found with Q44R/H45R, Q44R/H45R/M115K, but not M115K (Figure 2C). Thus, although in cells all three mutants with the Q44R/H45R change can form dimers, they do not all appear to be stable *in vitro* (Figure 2A-B). Collectively, the data suggested that amino acids 44 and 45 are key for rhA3C dimerization, but there may be different amino acids required to stabilize rhA3C dimerization than for hA3C.

### Simulated introduction of rhA3C amino acids into hA3C disrupts dimerization

The prediction from the SEC data is that introduction of the rhA3C amino acids into hA3C would disrupt hA3C dimerization. Biochemical analysis of the dimeric hA3C S188I and cA3C demonstrated that the dimer in the hA3C crystal structure used the same interface as these natural dimeric forms of A3C (Adolph, Ara, et al., 2017; Kitamura et al., 2012). Thus, MD simulations of hA3C were performed to test the effects of rhA3C amino acids on dimerization. The single amino acid changes converting hA3C amino acids to those of rhA3C, N115M, R44Q, and R45H were tested alone and in combination.

The hA3C R44Q/R45H/N115M mutant showed only slight changes in root mean squared fluctuation (RMSF), with 20 amino acids across both monomers exhibiting a change greater than 0.5 Å, and only 4 amino acids changing by more than 1.0 Å (Figure 3A). These changes were scattered in both increases and decreases at adjacent residues, with very few trends spanning any region of the protein. This low degree of impact to the dynamic motion is consistent with the negligible (less than 10%) change in the intermolecular hydrogen bonds (Figure 3B). By comparison, positions 44 and 45 have five interactions decreasing by more than 10% and one increasing by more than 10% (Figure 3B). Normal mode analysis based on the trajectories indicated that the essential dynamics of all the systems are captured by the first two modes (Figures S2 and S3). Difference correlation and principal component analysis (PCA) comparing the first two modes of motion show the hA3C R44Q/R45H/N115M variant to be qualitatively similar to the hA3C (Figure 3C-D). However, energy decomposition analysis indicates that the interaction energy between the two monomers is destabilized by approximately 630 kcal/mol with respect to the hA3C (Table S1). This destabilization is similar to that of the hA3C R44Q/R45H double mutant, suggesting that hA3C N115M has an effect if in the context of a R44Q/R45H mutant (Table S1).

**Figure 3.**
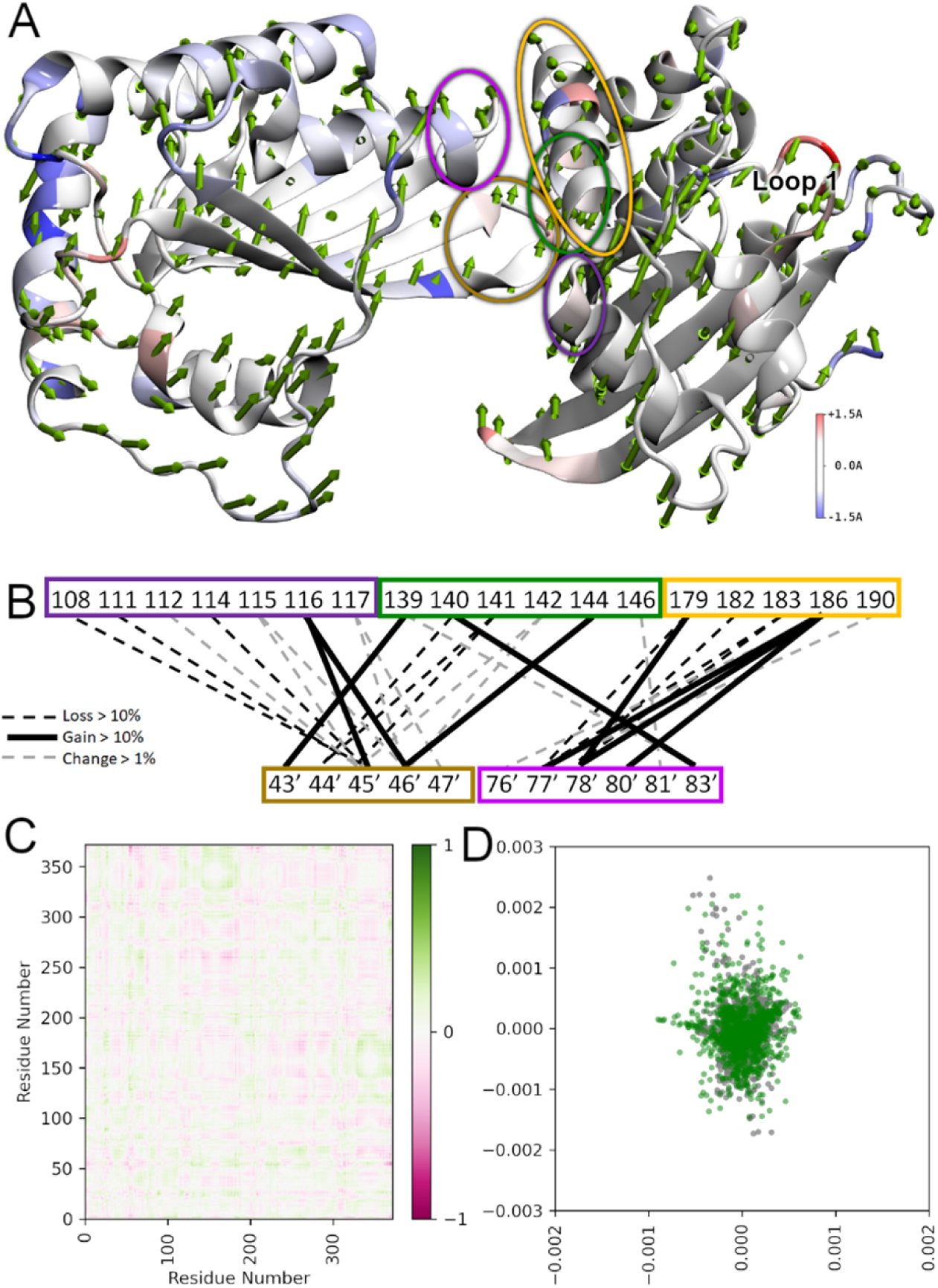
Model analysis of hA3C R44Q/R45H/N115M illustrates changes from hA3C WT. The hA3C R44Q/R45H/N115M variant compared to hA3C WT. **(A)** RMSF heatmapped to protein structure with first normal mode shown as arrows from each amino acid in sequence. Arrow direction and magnitude shows the motion of each residue as a portion of the total largest contributor to essential motion. Arrows of larger magnitude contribute more to that essential motion. The RMSF heatmapping shows the total fluctuation of each residue from an average position over the trajectory. Higher values of RMSF indicate greater movement from this average position. Loop 1 that is important for activity is labeled. **(B)** Dimer interface hydrogen bonding changes. Colored blocks of residues correspond to regions encircled in panel (A) with the same color. Black hatched lines mean a loss >10%, black lines mean a gain of >10%, and grey hatched lines mean a change of >1%. **(C)** Difference correlation plot between hA3C R44Q/R45H/N115M and hA3C WT. **(D)** The PCA showing first two modes of hA3C R44Q/R45H/N115M (green) against hA3C WT (grey). Each dot represents an amino acid.

We analyzed the single mutants to determine if they showed a destabilizing effect compared to the triple mutant, hA3C R44Q/R45H/N115M. Analyzing the essential dynamics modes using PCA it was found that R45H showed minimal qualitative changes in dynamic motion compared with hA3C, N115M, and R44Q (Figure S4). The N115M variant showed some change in the first normal mode, and R44Q, showed greater scatter on the first two modes (Figure S4). These changes correspond to the largest changes in intermolecular hydrogen bonds as well, suggesting that the R44Q and N115M mutations have a noticeable impact on dynamic motion and dimer interface contacts. Correlated motion across all single mutants was qualitatively similar compared to the hA3C WT, again with the exception of N115M, which showed greater patterns of correlated motion (Figure S5). Additionally, dimer interaction energies were shown to be destabilized in R44Q, R45H, and N115M single mutants, compared with hA3C WT (Table S1).

The hA3C R44Q/R45H double mutant system showed greater destabilization based on the MD simulations results compared with the sum of the two single mutants R44Q and R45H, suggesting that either of the arginines at these locations may compensate for the loss of the other, but the loss of both is significant (Table S1). Altogether, these results show significant disruptions in dimer interface hydrogen bonding when hA3C amino acids are converted to rhA3C amino acids. This is consistent with the SEC data, which showed that the reverse changes, rhA3C amino acids converted to hA3C amino acids promote dimerization (Figure 2A-B) and place a significant role on amino acids 44 and 45.

### rhA3C deaminase activity is increased with hominid-like amino acids at positions 44 and 45

Since the rhA3C appeared to have different determinants for stable dimerization than hA3C, we wanted to test that dimerization did indeed correlate with increased deamination activity in rhA3C. We examined the specific activity of rhA3C by conducting a uracil DNA glycosylase assay with a 118 nt substrate containing two 5’TTC motifs (Figure 4A-B). We used a time course of deamination to determine the linear range of activity (Figure 4A-B) and then calculated the specific activity of the rhA3C in that region (i.e., 5 to 15 min). We found that the rhA3C WT was approximately 2-fold less active than the three rhA3C mutants containing the Q44R/H45R change, confirming that residues that promote dimerization are of primary importance to rhA3C activity (Figure 4C). However, since the rhA3C needs to search the DNA for the 5’TTC motifs among the non-motif containing DNA, the specific activity is made up of both the chemistry in the catalytic site once reaching the substrate C and the time it took to search for the 5’TTC motif. The search occurs by facilitated diffusion and results in the enzyme being processive, i.e., deaminating more than one 5’TTC motif in a single enzyme substrate encounter. The processivity was calculated by analyzing the individual bands from the uracil DNA glycosylase assay (Figure 4B). The processivity requires single-hit conditions in which an enzyme would have acted only on one DNA (see Materials and Methods).

**Figure 4.**
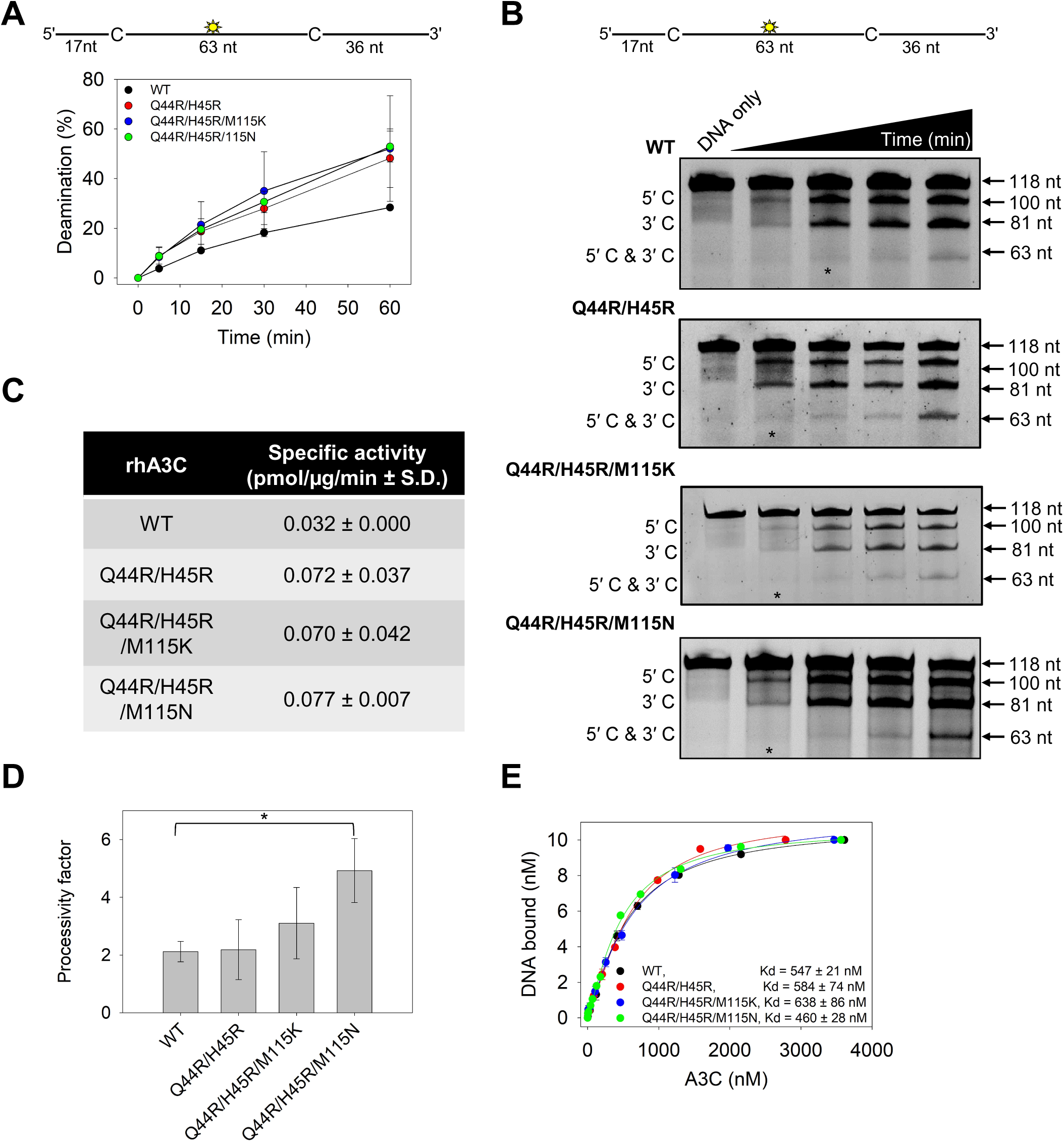
Arginines at positions 44 and 45 in rhA3C increase enzyme activity. (**A-B**) Time course of rhA3C WT, Q44R/H45R, Q44R/H45R/M115K, and Q44R/H45R/M115N on a 118 nt fluorescently labeled ssDNA with two 5′TTC deamination motifs spaced 63 nt apart. Reactions were performed with 100 nM substrate DNA and 1000 nM rhA3C for the indicated amount of time (5–60 min). **(B)** Representative gel from three independent deamination assay experiments. Lanes with an asterisk were used for processivity calculations shown in panel **(D)**. **(C)** Specific activity as calculated from the time course in **(A-B)** and showed increased activity if the rhA3C had the Q44R/H45R mutation. **(D)** Processivity of rhA3C WT, Q44R/H45R, Q44R/H45R/M115K, and Q44R/H45R/M115N calculated from the time course in **(A-B)**. The processivity factor measures the likelihood of a processive deamination over a nonprocessive deamination and is described further in the ‘Materials and Methods’ section. Designations for significant difference of values were \*\*\**P* ≤ 0.001, \*\**P* ≤ 0.01 or \**P* ≤ 0.05. **(E)** The apparent Kd of rhA3C enzymes from a 118 nt fluorescently labeled ssDNA was analyzed by steady-state rotational anisotropy for rhA3C WT, Q44R/H45R, Q44R/H45R/M115K, and Q44R/H45R/M115N. Apparent *K*_d_ values are shown in the figure with the standard deviation. **(A, D, E)** On the graphs, the standard deviation for three independent experiments is shown as error bars.

Using these conditions, we determined the number of deaminations at both 5’TTC motifs (5’C & 3’C) compared to the single deaminations at either the 5’TTC motif closest to the 5’-end (5’C) or 3’-end (3’C). From these values a processivity factor can be calculated which is the fold likelihood that a processive deamination would take place. We found that the processivity of rhA3C WT was 2.1 (Figure 4D). In comparison, a nonprocessive enzyme has a processivity factor of 1. Thus, the rhA3C WT is not very processive, consistent with it being a monomer. The mutants had small but consistent increases in processivity where the Q44R/H45R processivity factor was 2.5, the Q44R/H45R/M115K slightly more processive (processivity factor of 3.2) and the Q44R/H45R/M115N that could partially dimerize most stably had a 2-fold higher processivity factor than the WT (processivity factor of 4.9, Figure 4D). Consistent with the most stable dimerization, only the Q44R/H45R/M115N mutant had processivity increase that was significantly higher than the WT (Figure 4D). However, increases in processivity and specific activity did not increase the affinity of the A3C for the 118 nt ssDNA, as measured by steady-state rotational anisotropy (Figure 4E).

### Increase in rhA3C specific activity correlates with increase in HIV-1 restriction activity

To determine if the *in vitro* differences of rhA3C from hA3C affect HIV-1 ΔVif ΔEnv (referred to as HIV) restriction ability we used a single-cycle infectivity assay. We compared hA3C S188I that is restrictive to HIV and rhA3C. We found that hA3C S188I could restrict HIV, but rhA3C could not, despite naturally having amino acid I188 (Figure 5A). We also checked rhA3C activity against the OWM SIV from sooty mangabey monkey (smm). The rhA3C only restricted SIVsmm ΔVif 2-fold, in contrast to hA3C S188I (13-fold), hA3C (6-fold) and cA3C (4-fold) that were more restrictive (Figure S7). Notably, the hA3C S188I was able to restrict SIV ΔVif better than HIV, suggesting that SIV is restricted more easily (Figure 5A and Figure S7) (Adolph, Ara, et al., 2017). Thus, overall, the data show that rhA3C is not active against HIV or SIV.

**Figure 5.**
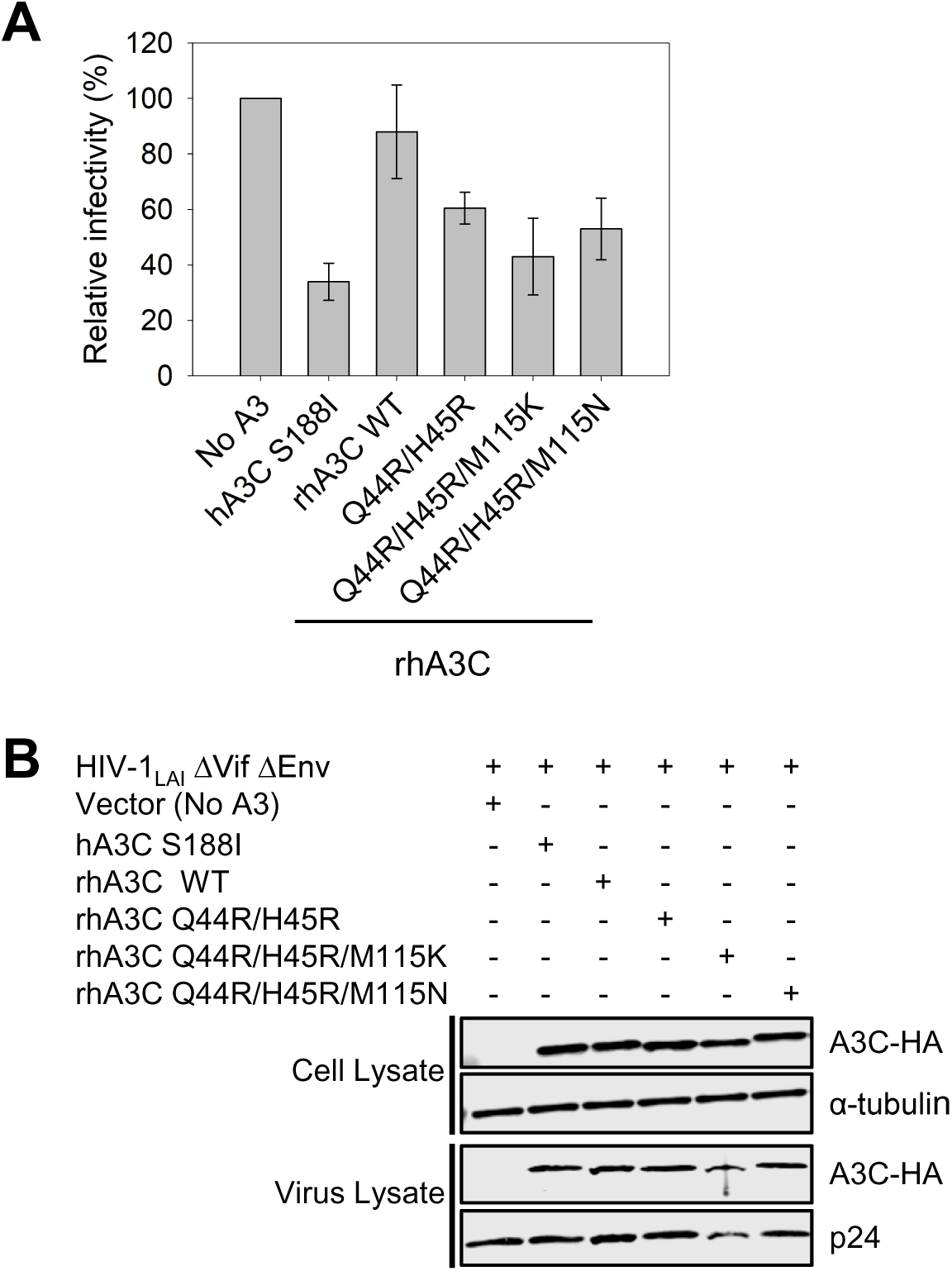
Amino acids 44 and 45 in rhA3C are key determinants for HIV restriction ability. **(A)** Infectivity was measured by β-galactosidase expression driven by the HIV-1 5΄LTR from TZM-bl cells infected with VSV-G pseudotyped HIV ΔVif ΔEnv that was produced in the absence or presence of 3xHA tagged hA3C S188I, rhA3C WT, and rhA3C mutants Q44R/H45R, Q44R/H45R/M115K, and Q44R/H45R/M115N. Results normalized to the no A3 condition are shown with error bars representing the Standard Deviation of the mean calculated from three independent experiments. (**B**) Immunoblotting for the HA tag was used to detect A3C enzymes expressed in cells and encapsidated into HIV ΔVif ΔEnv pseudotyped virions. The cell lysate and virion loading controls were α-tubulin and p24, respectively.

To determine if the lack of rhA3C restriction was due to it being a monomer, we tested the rhA3C dimer interface mutants. We began by converting only the rhA3C Q44/H45 to R44/R45 (as in hA3C). We found that this mutation alone resulted in a 1.5-fold increase in restriction from rhA3C wild type (WT) (Figure 5A). The WT and mutant were expressed and encapsidated into virions similarly (Figure 5B). Upon the addition of M115K and M115N mutations in a Q44R/H45R background, we found that there was a small increase in restriction. These triple mutants had a 1.7- to 2- fold increase in restriction activity against HIV in comparison to rhA3C WT, which was not due to increased encapsidation into virions (Figure 5B). Importantly, just the changes at amino acids 44 and 45 increased HIV restriction similarly to changes at position 115 (Figure 2A-B). Altogether, the data showed that the 2-fold increase in specific activity (Figure 4C) corresponded well with an approximate 2-fold increase in virus restriction (Figure 5C) and suggested that processivity may not be essential to HIV restriction by rhA3C. Further, these data demonstrated that rhA3C restriction activity was improved by converting dimer interface amino acids to the hA3C or cA3C amino acids, demonstrating that the rhA3C WT is a monomer form is less able to restrict HIV. However, the restrictive activity of the triple mutants was still 2-fold lower than that of hA3C S188I (Figure 5A) which suggested that there were likely additional determinants for dimerization in rhA3C.

### A key residue for rhA3C dimerization is uncovered via direct coupling analysis (DCA)

Since we could not achieve stable dimerization or HIV restriction activity equivalent to hA3C S188I from mutation of rhA3C amino acids 44/45/115 we turned to DCA, also termed coevolutionary analysis (Morcos et al., 2011). Since dimerization is relevant for catalytic activity, we hypothesized that amino acid interactions that maintain function must be encoded in the evolutionary history of the A3 family of sequences (Pfam PF18771). The sequences included any organism and homologs that have a sequence that is classified as a member of the A3 family. A hidden Markov Model profile (HMMER) was used to search the NCBI database, including non-curated sequences, to find sequences whose statistics look like members of the family. This allowed inclusion of more distant sequences as opposed to direct matches to A3C. A resulting multiple sequence alignment (MSA) with ∼2500 sequences was compiled (see Materials and Methods and Supplementary File 1). Methods like coevolutionary analysis have been useful to identify amino acid coevolution in sequence alignments and to predict residue-residue contacts in the 3D structure of proteins (Anishchenko, Ovchinnikov, Kamisetty, & Baker, 2017; Morcos et al., 2011) and to predict 3D folds (Sulkowska, Morcos, Weigt, Hwa, & Onuchic, 2012:Marks, 2011 #1265).

Interestingly, coupled amino acid changes that preserve dimeric interactions at the interface between oligomers can also be inferred and used to predict complex formation in dimers (dos Santos, Morcos, Jana, Andricopulo, & Onuchic, 2015; Malinverni, Jost Lopez, De Los Rios, Hummer, & Barducci, 2017; Ovchinnikov, Kamisetty, & Baker, 2014). Therefore, we analyzed an alignment of approximately 2500 sequences (see Materials and Methods) to identify the most important dimeric coevolved interactions for the A3 family. Using DCA we identified a set of residue-residue pairs that are important for dimerization (Figure 6 and Materials and Methods). Figure 7A shows, in a contact map, the residue-residue contacts found in the crystal structure of hA3C (PDB 3VOW) for monomeric interactions (light gray) and the dimeric contacts (blue). In the same map, red dots indicate the top 300 coevolving interactions found by DCA. We notice that monomeric contacts can be predicted from the analysis of sequences, but more relevant to our study, we uncover a region (Figure 7B) that coincides with homodimeric contacts in the crystal structure (Figure 7C).

**Figure 6.**
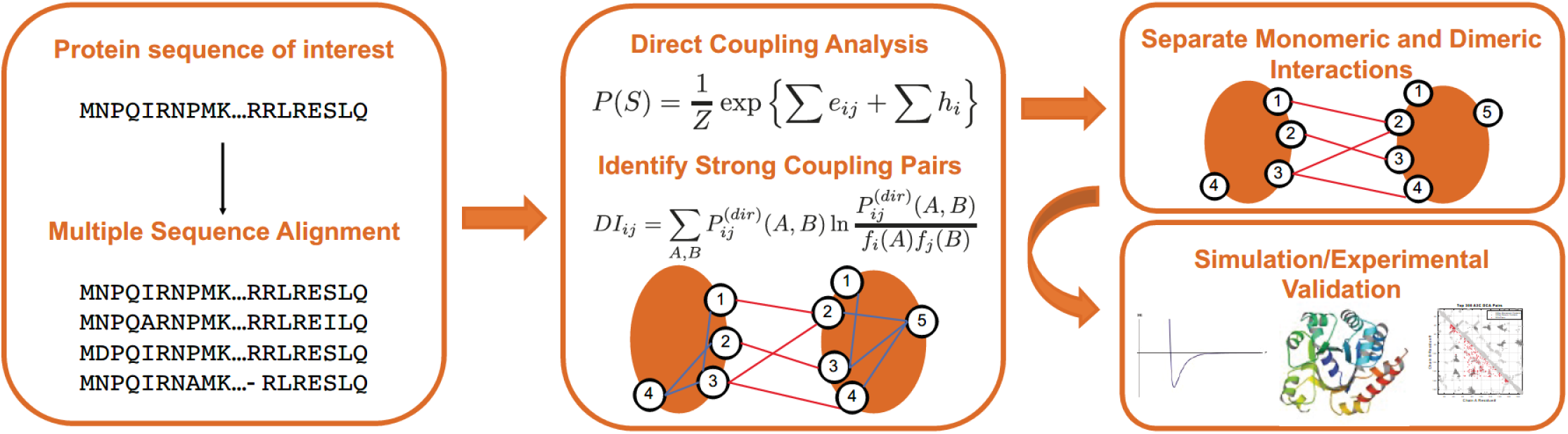
Coevolutionary analysis to identify relevant interaction residues involved in dimerization. First a multiple sequence alignment (MSA) is created to identify members of the A3C family. The MSA is then processed using Direct Coupling Analysis and a metric of coupling strength called Direct Information (DI). Once the top DI pairs have been identified, the monomeric crystal structure is used to distinguish monomeric from dimeric interactions. The resulting interacting residues are used to drive a simulation that predicts dimeric complexes. The residues involved in more coupled interactions are then proposed for experimental validation.

**Figure 7.**
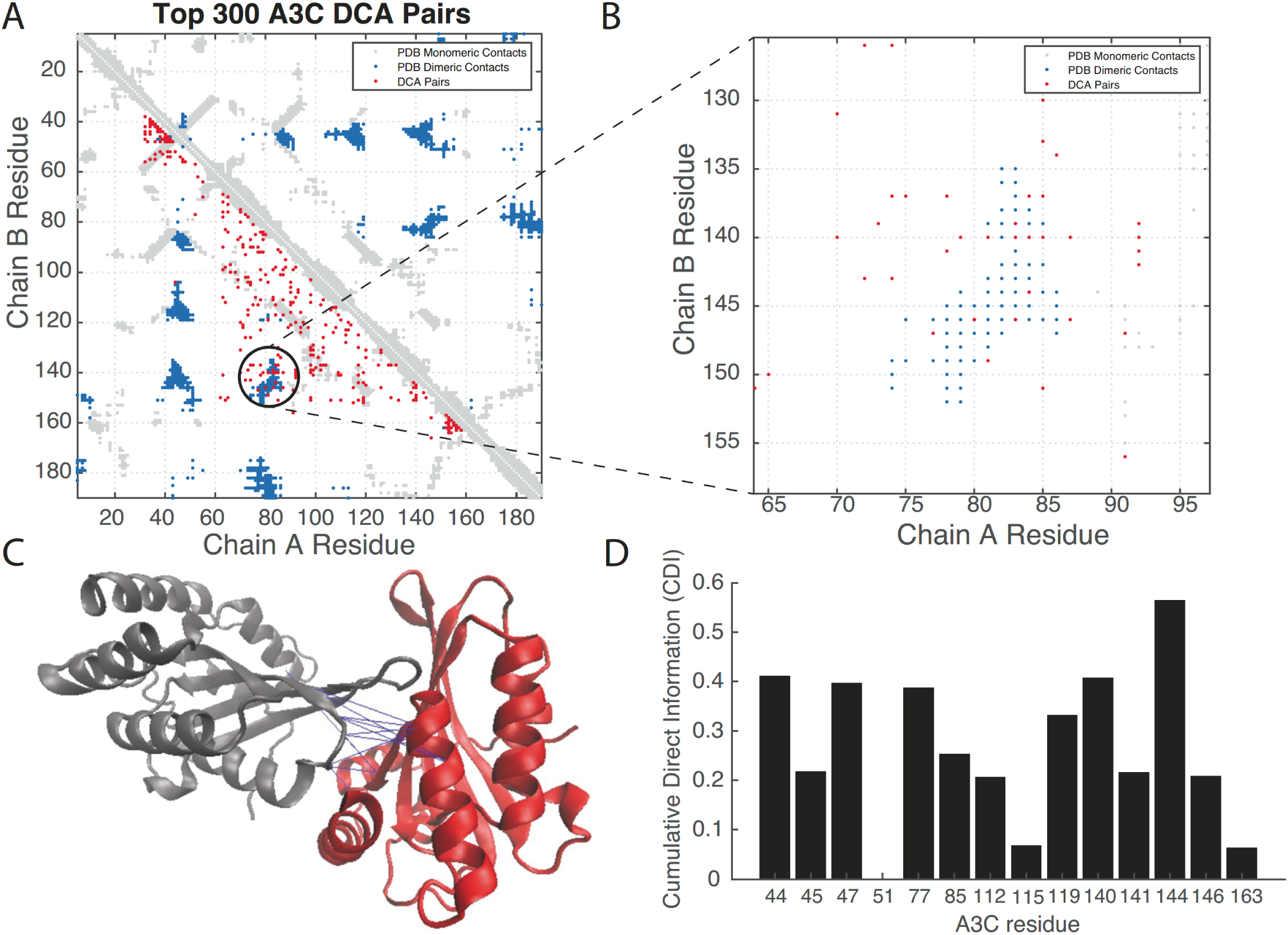
Directly coupled residue pairs identify relevant dimeric interacting residues preserved through evolution. **(A)** A contact map comparing the residue contacts in the x-ray crystal structure (3VOW) of hA3C (light gray for monomeric, dark blue for dimeric) and those interactions found by DCA (red). **(B)** A region of dimeric interactions shared by the crystal and coevolutionary analysis, highlighting the importance of those residues for dimer formation. **(C)** Overlay of the coevolved pairs on the 3VOW structure, depicting pairs inferred by the coevolutionary analysis. **(D)** A Cumulative Direct Information (CDI) metric identifies relevant residues at the interface. Of note, is the residue 144 that appears to have strong interactions with several residues and therefore a candidate for mutational analysis.

To further validate the relevance of those contacts for homodimeric interactions, we ran a coarse-grained MD simulation that uses such coupled coevolved pairs to drive a dimerization process (see Materials and Methods). The outcome of such simulation is a homodimer predicted completely from coevolutionary signatures. We notice that this predicted complex is not identical to the x-ray structure, but it does share a large portion of the homodimeric interface (Figure S8).

Having identified an evolutionarily relevant interface for dimerization, we proposed a metric to identify key residues for dimer formation and that at the same time are part of the set of amino acid differences between hA3C and rhA3C. Our metric, called Cumulative Direct Information (CDI) quantifies how much a given residue is involved in coevolving interactions from the most important ranked residue pairs. We reasoned that if a residue is involved in several coevolving interactions, then they could be ideal candidates for mutational studies. Figure 7D, shows this quantification for the top 200 dimeric interactions in the A3 family and for the residues that differ among the hA3C and rhA3C. We first noticed that the area between residues 44-47 was important according to this quantification, which was consistent with the experimental data (Figure 2) and MD simulations (Figure 3). This analysis showed that residue 115 does not have a high CDI score, which agrees with our results of a limited impact of residue 115 on activity beyond residues 44 and 45 (Figure 4 and Figure 5). More importantly, Figure 7D shows a dominant role of residue 144 in dimerization. Therefore, predicting 144 as a potentially important candidate for further mutational analysis.

### MD simulation predicts an important role for amino acid 144 in dimerization

Working with the hA3C crystal structure to convert amino acids to the rhA3C amino acids we tested if amino acid 144 influenced hA3C dimerization. The hA3C A144 was changed to a rhA3C S144 in the context of changes at amino acids 44 and 45, forming hA3C R44Q/R45H/A144S (rhA3C-like). This mutant exhibited greater than 0.5 Å change in RMSF on 119 amino acids, and 9 changing by more than 1.0 Å with respect to the hA3C WT (Figure 8A). This corresponded to larger regions of change on the protein, most particularly on loop 1, which is located between α-helix 1 and β-strand 1 (Figure 1 and Figure 8). PCA revealed a clear change in the dynamic motion especially on the first mode (Figures S2-S4). These larger changes are observed alongside significant loss of hydrogen bonding at the dimer interface by 0.7 time-averaged hydrogen bond (Figure 8B). The hydrogen bonds at the three mutation sites all showed considerable change, with position 144 increasing by more than 10% and positions 44 and 45 having six interactions decreasing by more than 10% (Figure 8B). The correlated motion of the variant is noticeably different compared with the hA3C, with a general trend of loss of correlated motion between the two monomers, consistent with a loss of dimer character (Figure 8C and Figure S4). These changes in correlated motion are also consistent with the observed changes in RMSF (Figure S6). Energy decomposition analysis indicates the dimer interaction is destabilized by approximately 770 kcal/mol (Table S1). This destabilization is greater than the sum of the individual variants, consistent with what has been previously observed with multiple mutants (Werner, Gapsys, & de Groot, 2021).

**Figure 8.**
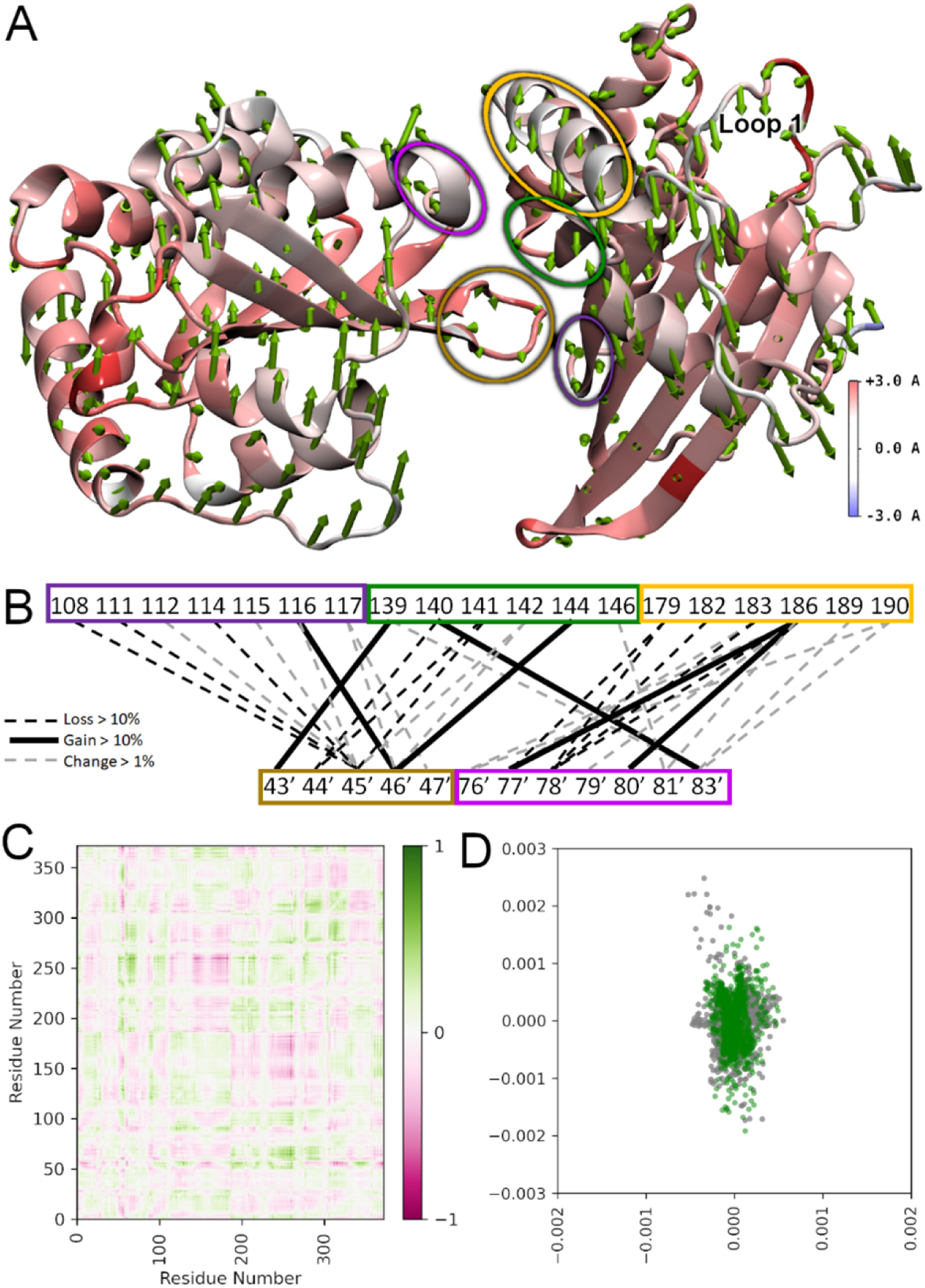
Model analysis of hA3C R44Q/R45H/A144S shows large changes in specific conformations. The hA3C R44Q/R45H/A144S variant compared to hA3C WT. **(A)** RMSF heatmapped to protein structure with first normal mode shown as arrows from each amino acid in sequence. Arrow direction shows the motion of each residue as a portion of the total largest contributor to essential motion. Arrow size denotes magnitude of motion. The RMSF heatmapping shows the total fluctuation of each residue from an average position over the trajectory. Higher values of RMSF indicate greater movement from this average position. Loop 1 that is important for activity is labeled. **(B)** Dimer interface hydrogen bonding changes. Colored blocks of residues correspond to regions encircled in panel (A) with same color. Black hatched lines mean a loss >10%, black lines mean a gain of >10%, and grey hatched lines mean a change of >1%. **(C)** Difference correlation plot between hA3C R44Q/R45H/A144A and hA3C WT. **(D)** PCA showing first two modes of hA3C R44Q/R45H/A144A (green) against hA3C WT (grey).

### A rhA3C S144A mutant in combination with Q44R/H45R enables stable dimerization and increased catalytic activity by a unique mechanism

Based on the predictions from coevolutionary analysis and MD simulations, we produced from *Sf*9 cells a rhA3C S144A mutant alone and in combination with changes at residues 44 and 45 to create a rhA3C Q44R/H45R/S144A mutant (hA3C-like). Consistent with the coevolutionary analysis, the S144A mutation enabled rhA3C to interact with itself (Figure 9A-B). However, a prominent dimer peak was only formed in the presence of a Q44R/H45R background (Figure 9A-B). Consistent with the contribution to activity of amino acids 44 and 45 the rhA3C Q44R/H45R/S144A mutant, but not the rhA3C S144A mutant alone, increased activity 2-fold compared to rhA3C WT (Figure 10A-C). Surprisingly, this increase in specific activity did not result in an increase in processivity (Figure 10D). There was also no change in ssDNA binding from rhA3C WT (Figure 10E).

**Figure 9.**
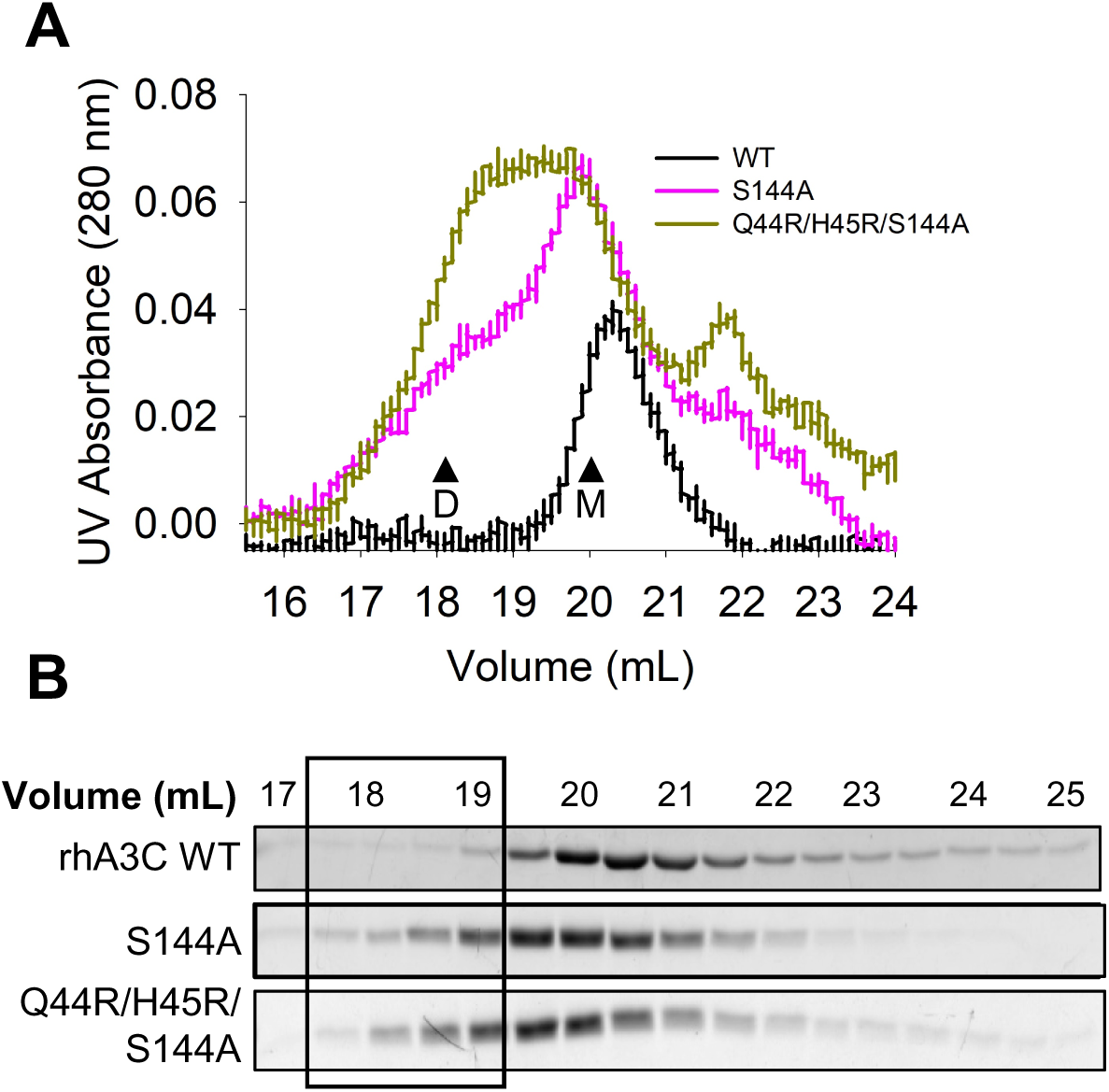
Mutation of rhA3C amino acids 44, 45, and 144 to hA3C amino acids enables dimerization. **(A)** SEC profile for rhA3C WT and rhA3C mutants S144A, and Q44R/H45R/S144A. Elution profiles for rhA3C WT was composed of a monomer peak (M, ∼20 ml elution volume). The rhA3C S144A showed a monomer and dimer peak (D, ∼19 mL elution volume). Only for the rhA3C Q44R/H45R/S144A there was a more prominent dimer peak. The elution profiles are shown as the UV absorbance during SEC elution. **(B)** Coomassie stained protein fraction resolved by SDS-PAGE that correspond the eluted fractions in **(A)**. Box shows where the dimeric fractions eluted.

**Figure 10.**
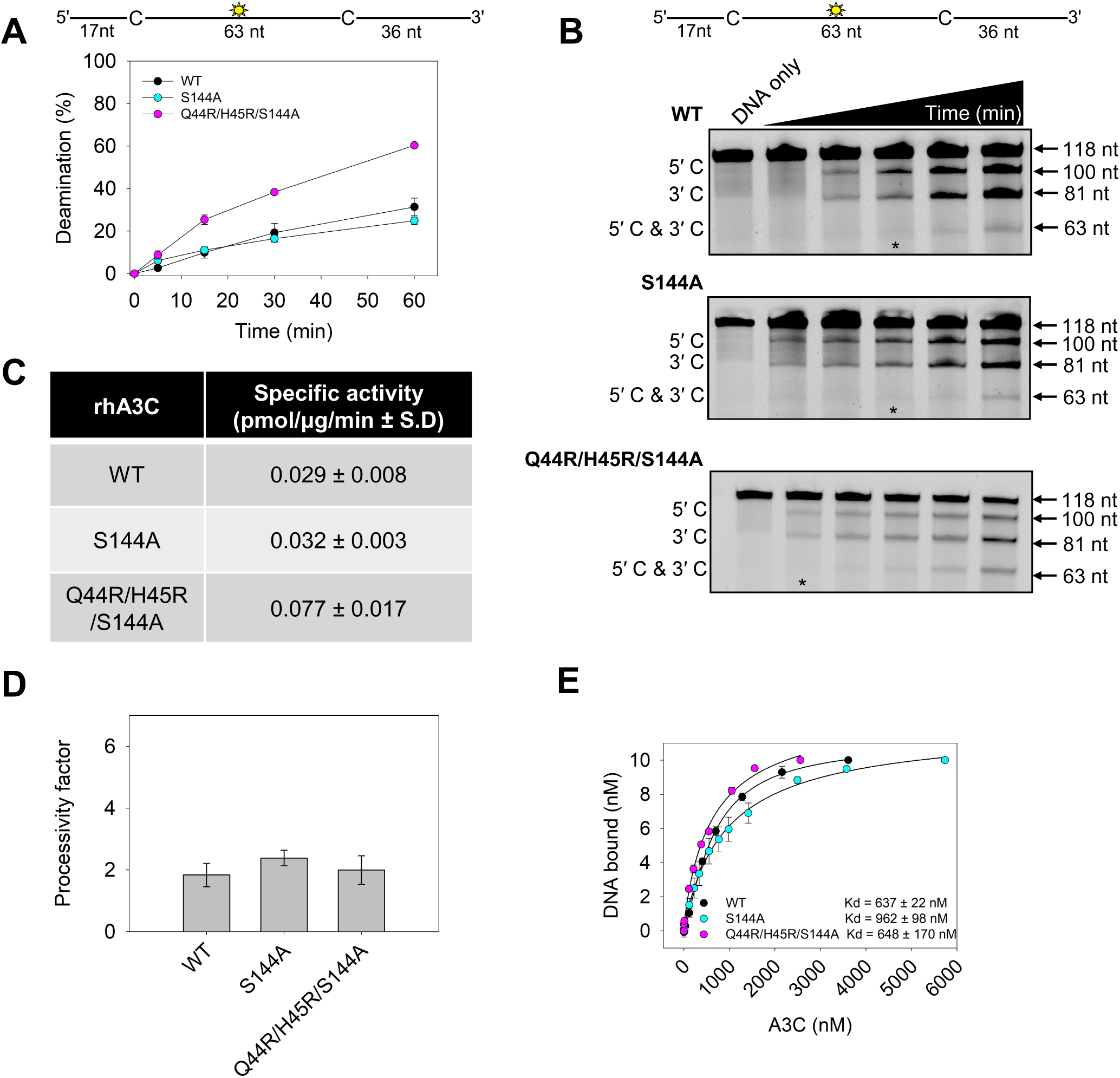
An S144A amino acid change in rhA3C increases deamination activity, but not processivity. (**A-B**) Time course of rhA3C S144A and Q44R/H45R/S144A on a 118 nt fluorescently labeled ssDNA with two 5′TTC deamination motifs spaced 63 nt apart. Reactions were performed with 100 nM substrate DNA and 1000 nM rhA3C for the indicated amount of time (5–60 min). **(B)** Representative gel from three independent deamination assay experiments. Lanes with an asterisk were used for processivity calculations shown in panel (D). **(C)** Specific activity was calculated from the time course in **(A)** and showed increased activity if the rhA3C had the Q44R/H45R/S144A mutation. **(D)** Processivity of rhA3C S144A and Q44R/H45R/S144A calculated from the time course in **(A-B)** demonstrated that it was not increased above WT rhA3C. The processivity factor measures the likelihood of a processive deamination over a nonprocessive deamination and is described further in the ‘Materials and Methods’ section. **(E)** The apparent Kd of rhA3C enzymes from a 118 nt fluorescently labeled ssDNA was analyzed by steady-state rotational anisotropy for rhA3C WT, S144A, and Q44R/H45R/S144A. Apparent Kd values are shown in the figure with the standard deviation. **(A, D, E)** On the graphs, the standard deviation for three independent experiments is shown as error bars.

The reason for this increased dimerization and catalytic activity without the expected increase in processivity appears to be due to changes in loop 1. In multiple A3s, including A3C, loop 1 has been found to mediate accessibility of the ssDNA substrate to the active site (Jaguva Vasudevan et al., 2020; King & Larijani, 2017; Shi, Demir, et al., 2017). The MD simulations revealed effects of changes at residue 144 on loop 1 that were unique from changes at residue 115 (Figure 3, Figure 8, Figure S6, and Figure S9). In the hA3C R44Q/R45H/A144S mutant, the increased RMSF on loop 1 corresponds to a reduction in dynamic motion compared with hA3C WT (Figure 8A and Figure S9). The helices around the active site maintain similar RMSF across the hA3C WT and hA3C R44Q/R45H/A144S mutant consistent with a specific change in loop 1 mediating the changes in specific activity (Figure 8). In the rhA3C, the opposite effect is expected, where the dynamic motion of rhA3C Q44R/H45R/S144A is increased compared to rhA3C WT. This is consistent with rhA3C Q44R/H45R/S144A having more time in a state with an open loop 1, which has been linked to increases in deamination activity and is consistent with our observation of increased specific activity (Figure 8, Figure 10C, and Figure S9).

#### Efficient HIV restriction by rhA3C when dimerization is mediated by amino acid 144

To test if the rhA3C S144A-mediated dimer form enabled HIV restriction, we conducted a single-cycle infectivity assay. We found that the rhA3C Q44R/H45R/S144A (Figure 11A, 33% infectivity) could restrict HIV at an equivalent level to hA3C S188I (Figure 11A, 24% infectivity). This was not due to increased encapsidation as it was encapsidated into HIV virions at an equivalent or slightly lesser amount that hA3C S188I (Figure 11B). Although the rhA3C S144A could dimerize, it could not restrict HIV, consistent with no increase in catalytic activity (Figure 9, Figure 10C, and Figure 11A). This result was interesting since both the S144A and M115N induced dimerization in combination with the Q44R/H45R mutations, but the rhA3C Q44R/H45R/S144A had a higher level of HIV restriction activity (Figure 10A, 33% infectivity and Figure 2B, 53% infectivity). We hypothesized that this simply could be because the Q44R/H45R/S144A had a more dominant dimer peak than Q44R/H45R/M115N (compare Figure 2A-B and Figure 9A-B). Alternatively, since the *in vitro* specific activities were similar, the difference could be due to a larger change of loop 1 for the 144 amino acid compared to 115 (Figure 8 and Figure S9). Since the rhA3C Q44R/H45R/S144A processivity was less than rhA3C Q44R/H45R/M115N (compare Figure 4D and Figure 10D). This led us to hypothesize that the rhA3C Q44R/H45R/S144A restricts HIV by a predominantly deamination-independent mode (mediated by nucleic acid binding) and rhA3C Q44R/H45R/M115N a deamination-dependent mode (mediated by deamination).

**Figure 11.**
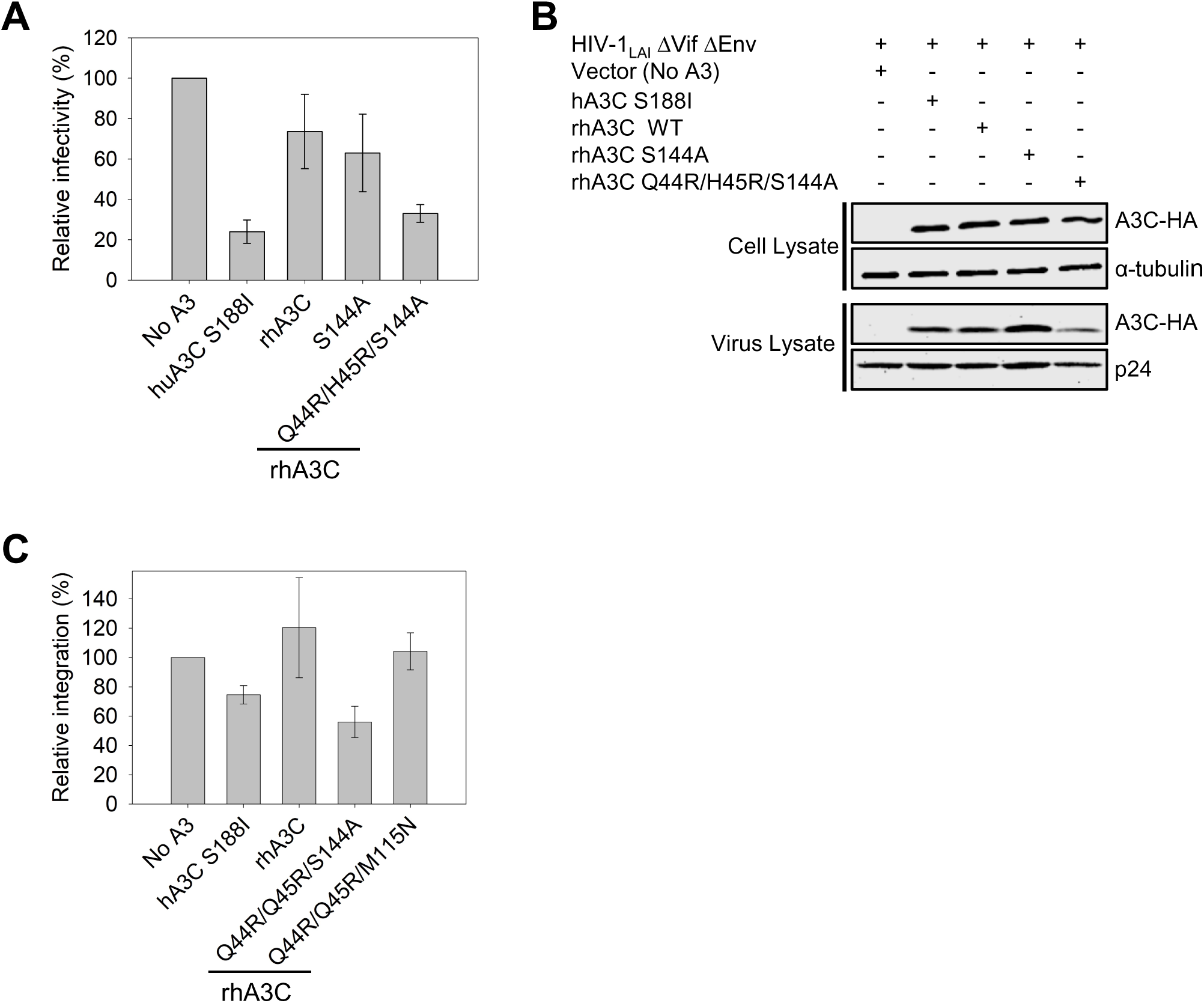
Amino acids 144 in combination with 44 and 45 in rhA3C enable HIV restriction ability. **(A)** Infectivity was measured by β-galactosidase expression driven by the HIV-1 5΄LTR from TZM-bl cells infected with VSV-G pseudotyped HIV ΔVif ΔEnv that was produced in the absence or presence of 3xHA tagged hA3C S188I, rhA3C WT, and rhA3C mutants S144A and Q44R/H45R/S144A. Results normalized to the no A3 condition are shown with error bars representing the Standard Deviation of the mean calculated from three independent experiments. Immunoblotting for the HA tag was used to detect A3C enzymes expressed in cells and encapsidated into HIV ΔVif ΔEnv pseudotyped virions. The cell lysate and virion loading controls were α-tubulin and p24, respectively. **(C)** The relative amount of proviral DNA integration in infected HEK293T cells in the presence of hA3C S188I, rhA3C WT, and rhA3C mutants Q44R/H45R/S144A and Q44R/H45R/M115N in comparison to the No A3 condition was determined by qPCR. Error bars represent the standard deviation of the mean calculated from at least two independent experiments.

We tested the deamination-dependent mode by determining the level of G→A mutations in coding strand of the integrated proviral DNA. We tested hA3C S188I, rhA3C WT and the two rhA3C mutants, Q44R/H45R/S144A and Q44R/H45R/M115N. The hA3C S188I had the highest level of G→A mutations (Table 1, 6.54 G→A mutations/kb). The majority mutations were in the GA→AA context (5’TC on the (-)DNA), which is the commonly preferred context for A3C (Table 1). Consistent with lower activity against HIV (Figure 11A), the rhA3C WT had the lowest level of G→A mutations (Table 1, 2.15 mutations/kb). For rhA3C WT there was an approximately equal amount of mutations with a GG→AG context (5’CC on the (-)DNA) and GA→AA context, indicating that the rhA3C active site has a more relaxed sequence preference. The GG→AG context is primarily associated with A3G (Yu et al., 2004). The rhA3C Q44R/H45R/S144A induced 1.5-fold less mutations than the rhA3C Q44R/H45R/M115N (Table 1, 3.20 and 4.67 G→A mutations/kb, respectively) consistent with rhA3C Q44R/H45R/S144A being 2-fold less processive than rhA3C Q44R/H45R/M115N.

**Table 1.** Analysis of A3-induced mutagenesis of *pol* gene from integrated HIVΔVif.

| **A3 enzyme** | **Base Pairs**<br>**Sequenced** | **Total**<br>**G→A**<br>**mutations** | **Total**<br>**GG→AG**<br>**mutations** | **Total**<br>**GA→AA**<br>**mutations** | **G→A**<br>**Mutations**<br>**per kb** | **GG→AG**<br>**mutations**<br>**per kb** | **GA→AA**<br>**mutations**<br>**per kb** |
| --- | --- | --- | --- | --- | --- | --- | --- |
| **No A3** | 8715 | 4 | 2 | 2 | 0.46 | 0.23 | 0.23 |
| **hA3C S188I** | 8715 | 57 | 15 | 41 | 6.54 | 1.72 | 4.70 |
| **rhA3C**<br>**WT** | 6972 | 15 | 6 | 7 | 2.15 | 0.86 | 1.00 |
| **rhA3C**<br>**Q44R/H45R**<br>**/S144A** | 8134 | 26 | 12 | 11 | 3.20 | 1.48 | 1.35 |
| **rhA3C**<br>**Q44R/H45R**<br>**/M115N** | 8134 | 38 | 20 | 15 | 4.67 | 2.46 | 1.84 |

To further explore how the rhA3C Q44R/H45R/S144A could restrict HIV more than rhA3C Q44R/H45R/M115N we also determined the level of proviral DNA integration. A3s can inhibit reverse transcriptase activity, which results in less completed proviral DNA synthesis, and less integration. The rhA3C WT and rhA3C Q44R/H45R/M115N did not decrease proviral DNA integration (Figure 11C). However, hA3C S188I and rhA3C Q44R/H45R/S144A did decrease proviral DNA integration (Figure 11C). The rhA3C Q44R/H45R/S144A allowed only 56% of the total proviral DNA to integrate, relative to the no A3 condition (Figure 11C). These data show that the rhA3C Q44R/H45R/S144A dimer restricts HIV using both deamination - dependent and -independent modes of restriction which are more effective than only the deamination-dependent mode used by rhA3C Q44R/H45R/M115N.

### Evolutionary dynamics of key residues involved in A3C dimerization

Because amino acids at residues 44, 45, 115, and 144 were all found to be important in the gain of dimerization and restriction activity of rhA3C, we examined the evolution of these residues over primate evolution. None of these residues correspond to those that were previously reported to be under positive selection (Wittkopp et al., 2016). Nonetheless, they do vary in Old World Primates and in Hominoids. While both the rhesus and the crab-eating macaque encode the QHMS at residues 44, 45, 115, and 144, respectively (Figure 12), the Northern pig tiled macaque encodes a cysteine (C) at residue 45, while the nearest outgroup, the Baboon, encodes an arginine (R) at residue 45, which is the residue found in hominids at that position (Figure 12). We used the sequences of 17 Old World Monkeys and Hominoids as well as one New World Monkey (for purposes of rooting the tree) to reconstruct the ancestral amino acids at each of these positions (Figure 12). We find evidence that the glutamine (Q) at amino acid 44 which is unfavorable for dimerization of rhA3C is, in fact, a derived trait as the ancestral amino acid at position 44 at both the root of the rhesus/baboon/drill common ancestor as well as the common ancestor that includes the african green monkeys is a histidine (H) at position 44. These results suggest that loss of dimerization of rhA3C is an ancient event, but may not have occurred by the same mechanism in all lineages.

**Figure 12.**
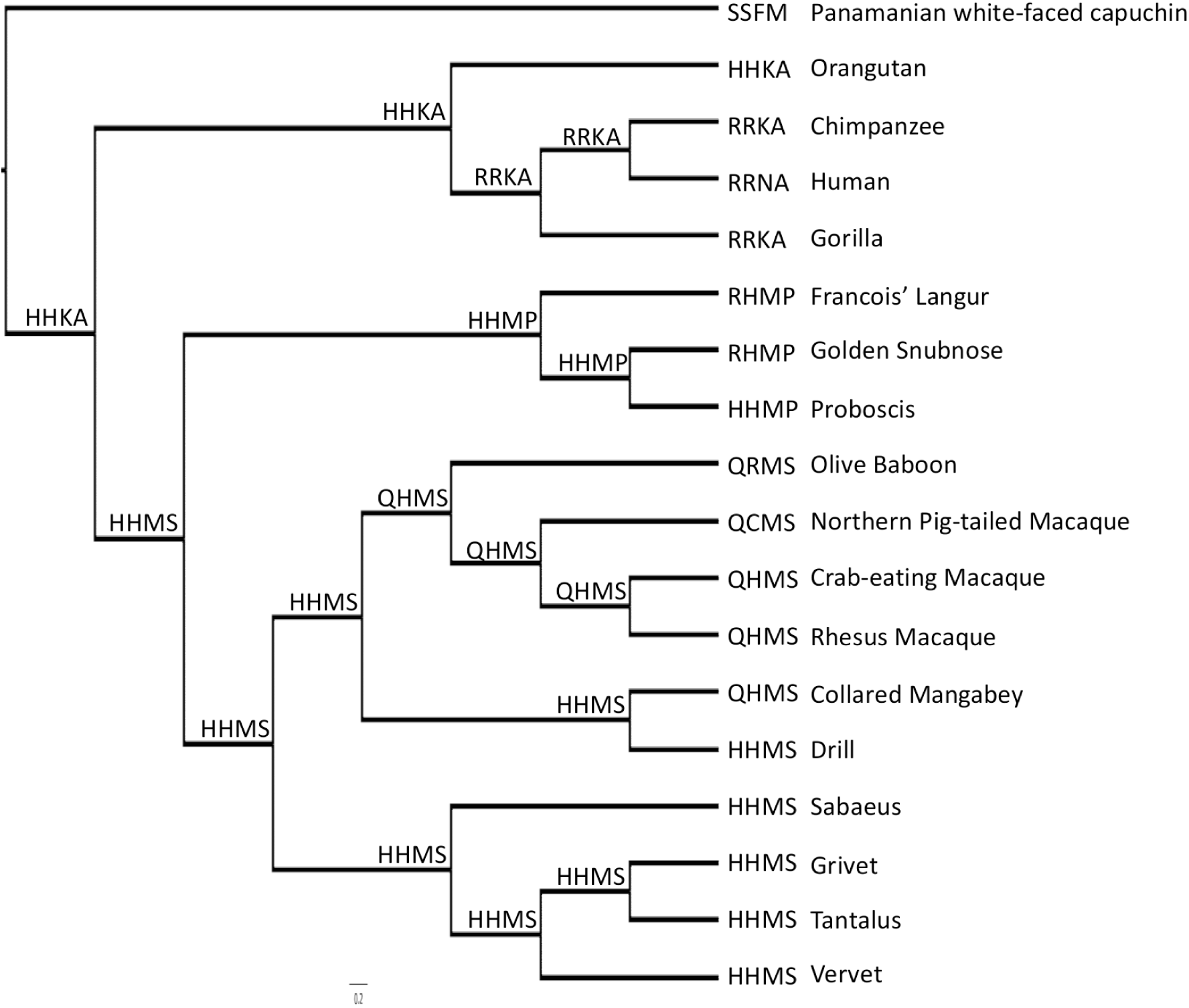
Ancestral reconstruction of residues involved in rhA3C dimerization. A phylogeny depicting residues crucial for A3C dimerization, including reconstructed ancestral residues. The species included are on the right of the phylogenetic tree of A3C sequences as well as the four amino acids at residues 44, 45, 115, and 144, respectively. The identities of the amino acids at each of these positions as calculated by FASTML are shown at each node of the tree.

## Discussion

Throughout the evolution of a host restriction factor, the protein must be able to retain activity against the virus, which may require compensatory mutations over time. In this study, we investigated the impact of evolutionary changes on a specific A3 enzyme, rhA3C. Interestingly, the majority of positive selection in A3C has taken place outside of the interaction motif with Vif, which is unique and suggests an evolution of enzyme activity, rather than avoidance of the viral antagonist, which provides a unique model to study restriction factor evolution (Wittkopp et al., 2016). Further, our work highlights the importance of an integrated strategy to identify relevant functional interactions in enzymes. In particular, it describes a cycle of experiment and theory that allowed us to efficiently identify key positions from a combinatorial web of potential interactions. This methodology is transferrable and warrants its application to study other members of the APOBEC family as our analysis showed that the evolutionary signals for dimerization seem to be preserved across multiple organisms and family members with distinct functions. This methodology would also be useful in other protein systems.

Determination of activity against lentiviruses for A3 enzymes is multifactorial. First, it requires viral encapsidation, which occurs for all A3C orthologues tested thus far (Figure 5) (Adolph, Ara, et al., 2017; Wittkopp et al., 2016). After that, there is a requirement to deaminate cytidines in lentiviral (-)DNA or inhibit reverse transcriptase (Harris & Dudley, 2015). Despite rhA3C inducing 2 mutations/kb, approximately 20 mutations in the total genome, the infectivity of the HIV was not significantly decreased (Figure 11 and Table 1). Since the mutations are stochastic, there may be some genomes with many mutations, some with none, and some with mutations that are not inactivating. Therefore, the more mutations, the better for ensuring inactivation. Increased processivity can achieve this, as evidenced by the 2-fold increase in mutations by the more processive rhA3C Q44R/H45R/M115N mutant, but this was still not enough for robust HIV restriction (Figure 5 and Table 1). What was needed was a deamination independent restriction of reverse transcriptase in combination with inducing mutagenesis to enable more robust restriction as for rhA3C Q44R/H45R/S144A (Figure 11 and Table 1).

However, the rhA3C Q44R/H45R/S144A results seemly go against numerous studies showing a direct correlation between processivity and mutation frequency in multiple A3 enzymes, including hominid A3Cs (Adolph, Ara, et al., 2017; Ara et al., 2014; Feng & Chelico, 2011; Feng et al., 2015). Using an *in vitro* system to study the effect of processivity on mutations during reverse transcription, it was shown that A3A, a non-processive enzyme, could introduce a similar number of mutations to A3G, a highly processive enzyme, but not at the same locations along the ssDNA (Love, Xu, & Chelico, 2012). The processivity was needed for mutations to accumulate in ssDNA regions rapidly lost to replication and dsDNA formation. The deaminations of A3A were achieved by a quasi-processive search involving multiple on and off interactions with the ssDNA. In this process time is lost, but deaminations still occur. This may be occurring with rhA3C Q44R/H45R/S144A.

Also interesting was that for rhA3C, the dimer did not always result in a processive enzyme. The rhA3C Q44R/H45R/S144A had equal specific activity to rhA3C Q44R/H45R/M115N but by a seemingly different mechanism than processivity (Figure 4 and Figure 10). In addition, the rhA3C S144A single mutant could dimerize, but had no increase in activity (Figure 9 and Figure 10). Where processivity makes the search for cytosines more efficient, the rhA3C Q44R/H45R/S144A is predicted to have an altered loop 1 that is able to increase the deamination activity (Figure 8 and Figure S9). Loop 1 has been identified to be a gate-like structure that controls access of the ssDNA to the active site (King & Larijani, 2017; Shi, Demir, et al., 2017). A3A that has an open loop 1 conformation, has high specific activity with low processivity (Shi, Carpenter, et al., 2017). In contrast, A3B and a related family member Activation Induced Cytidine deaminase (AID) have closed loop structures (King & Larijani, 2017; Shi, Demir, et al., 2017). However, A3B and AID are processive and still achieve similar activity to A3A, but are more selective for which cytidines are deaminated or under which types of conditions, e.g., A3B is more active when ssDNA is in excess to the enzyme (Adolph, Love, Feng, & Chelico, 2017). Our data suggest that the rhA3C Q44R/H45R/S144A has increased rhA3C activity in an A3A-like manner (Figure 10 and Figure 11). Further, the rhA3C S144A mutant demonstrates that dimerization was necessary, but not sufficient, since arginines at residues 44 and 45 were also required for the increase in specific activity and restriction activity (Figure 10 and Figure 11). The reason for why the processive rhA3C Q44R/H45R/M115N does not increase activity enough may simply be due to the unstable nature of the dimer, which would be needed in the absence of the loop 1 alteration (Figure 2). Loop 1 has previously been shown to regulate activity of hA3C (Jaguva Vasudevan et al., 2020). The hA3C was shown to have an amino acid pair 25W/26E that decreased activity compared to a hA3C 25R/26K mutant (Jaguva Vasudevan et al., 2020). Our MD simulations data suggest that alterations in the protein structure from a 144A substitution changed the dynamics of loop 1 indirectly, which also resulted in increased specific activity, in a 44R/45R background (Figure 8, Figure 10, and Figure S9).

Having determined the residues needed for activity against HIV in rhA3C we examined other OWMs. We found that none of the analyzed OWM A3Cs contained the right amino acid combinations for robust activity, although there were considerable changes. The rhA3C sequences at the four key amino acids is ancient, but is a derived trait from the common ancestor of Old World Monkeys (Figure 12). Other OWMs also had other changes to one or two sites. This perhaps indicates that different selective pressures were on OWM A3C from a non-lentivirus pathogen. This may be a retroelement, such as LINE-1, since hA3C can restrict LINE-1 similarly to hA3C S188I, indicating that there is no requirement for dimerization (Wittkopp et al., 2016). Alternatively, A3C may act in concert with other A3s. For example, in humans, A3G and A3F have been found to hetero-oligomerize and this increases the activity of both enzymes (Ara et al., 2017). This hetero-oligomerization has been understudied and perhaps A3C acts with another A3 and has greater activity. This would perhaps explain the different requirements for dimerization in comparison to hA3C although rhA3C and hA3C use the same general interface. Finally, it is possible that A3C, like A3H, has lost activity during evolution in primates (Garcia & Emerman, 2018), perhaps because of selection against the deleterious effects of the enzyme, or because its activity has been usurped by other A3 enzymes.

Altogether, the data show that rhA3C is not active against HIV, but activity can be enhanced through dimerization that either increases processivity or more robustly through dimerization that causes alteration of loop 1 conformation and dynamics. This is important for understanding the biochemical basis of activity of A3C and other A3 family members and for using it as a tool to predict anti-lentiviral activity in other OWMs. The DCA methodology could provide additional amino acids to test, however we tested the ones with the highest CDI score or importance based on hominid A3C. The OWM A3C has evidence of positive selection both within and outside the Vif binding region, suggesting that it has antiviral activity (Wittkopp et al., 2016). However, the fixation of the I188 in rhA3C and likely other OWMs did not impart anti-lentiviral activity, perhaps due to other compensatory mutations being made at the sites identified here, which were needed to combat other pathogens.

## Materials and Methods

### Plasmid constructs

The rhA3C DNA (GenBank: EU381233.1) was synthesized by GeneArt DNA synthesis and then subcloned into the appropriate vector. The rhA3C was cloned into the baculovirus transfer vector pFAST-bac1 containing an N-terminal GST tag as described previously using SmaI and NotI restriction sites (Feng et al., 2015), pcDNA3.1 vector with an C-terminal 3xHA tag using a XbaI restriction site, or pCMV vector with a N-terminal 3xFlag using NotI and SalI restriction sites. Site directed mutagenesis of rhA3C WT was used to create rhA3C M115K, rhA3C M115N, rhA3C Q44R/H45R, rhA3C Q44R/H45R/M115K, rhA3C Q44R/H45R/M115N, rhA3C S144A and rhA3C Q44R/H45R/S144A mutants. The hA3C, hA3C S188I, and cA3C have been previously described (Adolph, Ara, et al., 2017). All constructed plasmids were verified by DNA sequencing.

### Single-cycle infectivity assay

HEK293T cells (1 × 10^5^ cells per well) in 12 well plates were co-transfected with 500 ng of pHIV-1 LAI ΔVif ΔEnv, 180 ng of pMDG, which expresses VSV-G, and 100 ng of pcDNA A3C-3xHA expression plasmid using GeneJuice transfection reagent (EMD Millipore). The SIVsmm lineage 5 (L5) infectious molecular clone (Fischer et al., 2012) was obtained from Dr. Frank Kirchhoff with an inactivated Vif (SIVsmm ΔVif) (Nchioua et al., 2021). After 24 h post transfection the media was changed. After 44 h post transfection, culture supernatants containing the virus were harvested, filtered through 0.45 μm polyvinylidene difluoride (PVDF) syringe filters and used to infect TZM-bl cells. For infection of TZM-bl cells 1 x 10^4^ cells per well of a 96-well plate were infected with a serial dilution of virus in the presence of 8 μg/mL polybrene. Forty-eight hours after infection the cells were washed with PBS and infectivity was measured through colorimetric detection using a β-galactosidase assay reagent and spectrophotometer. Infectivity of each virus was compared by setting the infectivity of the “No A3” condition as 100%.

### Immunoblotting viral and cell lysates

A portion of the viral supernatant collected for infectivity assays was concentrated using the Retro-X (Clontech) following the manufacturer’s protocol. For immunoblotting 8 μL of concentrated virus was used. Producer cells were washed with PBS and lysed using 2× Laemmli buffer. Total protein in the cell lysate was estimated using the lowry assay and 30 µg total protein from each cell lysate was used for Western blotting. A3C was detected in cell lysates and virions using anti-HA antibody (Mouse monoclonal, Cat# H9658 (Sigma) for cell lysate, Rabbit polyclonal, Cat# H6908 (Sigma) for virus). Loading controls for cell lysates (α-tubulin, rabbit polyclonal, Cat # PA1-20988, Invitrogen) and virus (p24, mouse monoclonal, Cat #3537, NIH HIV Reagent Program) were detected using specific antibodies. Secondary detection was performed using Licor IRDye antibodies produced in goat (IRDye 680-labeled anti-rabbit 1: 10000 Cat # 926-68071 and IRDye 800-labeled anti mouse 1:10000 Cat # 926-32210).

### Proviral DNA sequencing

For proviral sequencing, 1 x10^5^ HEK293T cells per well of a 24-well plate were infected with supernatant containing virus in the presence of 8 μg/mL polybrene. The plates were spinoculated at 800 x *g* for 1 h. Cells were harvested after 48 h by removing the media, washing with PBS, and lysing the cells and extracting DNA with DNAzol (Invitrogen). The PCR amplification of a pol region of HIV-1 (581 bp) and treatment of DNA with DpnI was carried out as previously described (Ara et al., 2014). Primers have been previously described (Mohammadzadeh et al., 2019). Sequences were analyzed with Clustal Omega (Sievers et al., 2011) and Hypermut (Rose & Korber, 2000).

### Proviral DNA integration

Methods to quantify the integrated proviral DNA were adapted from (Belanger, Savoie, Rosales Gerpe, Couture, & Langlois, 2013). For infections, 1 × 10^5^ HEK293T cells per well of a 12-well plate were infected by spinoculation (1 h at 800* × g*) and in the presence of polybrene (8 μg/mL) with HIV produced from the single-cycle replication assays. DNA was extracted after 24 h using DNAzol (Invitrogen) according to manufacturer’s instructions. The DNA was then treated with DpnI and 50 ng was used in a PCR as previously described (Belanger et al., 2013). The PCR was then diluted 40-fold and used as the template in a qPCR as previously described (Belanger et al., 2013).

### Co-immunoprecipitation

For co-immunoprecipitation HEK293T cells in T75 cm^2^ flask were transfected with 1μg of each plasmid DNA using GeneJuice transfection reagent (EMD Millipore) as per manufacturer’s instructions. At 48 h post transfection, the cells were washed with PBS and lysed using IP buffer (50 mM Tris-Cl pH 7.4, 1% Nonidet-P40, 10% glycerol, 150 mM NaCl) supplemented with EDTA- free protease inhibitor (Roche). The protein concentration of cell lysates was measured using Bradford assay and equal amount of protein was added to anti-flag M2 magnetic beads (Sigma) in the presence of RNaseA (50 μg/ml; Roche) and incubated for 2 h with gentle rocking at 4°C. The beads were subsequently washed 5 times with the Tris-buffered Saline (TBS), and the immunoprecipitated proteins were subjected to SDS-PAGE and immunoblotting with anti-Flag antibody (Cat# F1804, Sigma), anti-HA antibody (Cat # H6908, Sigma). Secondary detection was performed using Licor IRDye antibodies produced in goat (IRDye 680-labeled anti-rabbit 1: 10000 Cat # 926-68071 and IRDye 800-labeled anti mouse 1:10000 Cat # 926-32210).

### Protein purification

The pFAST-bac1 vectors were used to produce recombinant baculovirus according to the protocol for the Bac-to-Bac system (Life Technologies), except using Insect GeneJuice (EMD Millipore) for bacmid transfection into *Sf*9 cells. The rhA3C WT and mutants were expressed in S*f*9 cells following infection with recombinant baculovirus with a multiplicity of infection of 2.5 and harvested after 72 hrs. The purification of cA3C has been previously described (Adolph, Ara, et al., 2017).

Pellets were stored at -80 °C until use. Thawed cell pellets from one litre cultures were resuspended in 35 ml of lysis buffer containing (20 mM HEPES (pH 7.5), 150 mM NaCl, 1% (v/v) Triton X-100, 10 mM NaF, 10 mM sodium phosphate, 10 mM sodium pyrophosphate, 100 µM ZnCl_2_, 1 mM EDTA, 10 mM DTT, 10% (v/v) glycerol and one Complete protease inhibitor tablet (Roche)). Following sonication and centrifugation, cleared lysate was incubated with glutathione-Sepharose resin (GE Healthcare) for several hours before washing the resin first with PBS buffer containing 1% (v/v) Triton X-100 and 500 mM NaCl followed by subsequent washes with PBS NaCl (250 mM). Finally, the resin was washed with digestion buffer containing 50 mM HEPES, pH 7.5, 250 mM NaCl, 1 mM DTT, and 10 % glycerol (v/v). The GST-A3C fusion enzyme containing slurry was treated with 0.02 units/μl thrombin (GE Healthcare) for a minimum of 4 hours at room temperature to release A3C from the GST tag. Purified proteins resolved by SDS-PAGE are shown in Figure S10.

### Size exclusion chromatography (SEC) and multi angle light scattering (MALS)

The SEC was performed using the Superdex 200 Increase 10/300 (GE Healthcare). Purified protein (230 ± 25 μg) was applied to the column equilibrated with 50 mM Tris (pH 8), 200 mM NaCl, 1 mM DTT. The flow rate was set to 0.6 ml/min and the eluate was monitored by the UV absorbance at 280 nm. Fractions were collected and further assessed using Coomassie stained SDS-PAGE. The same protocol was used to measure the light scattering and refractive index using a Wyatt Technology Multi-Angle Light Scattering (MALS) DAWN HELEOS II and Refractive Index (RI) OPTILAB T-rex, connected in tandem to Bio-Rad FPLC system.

### Deamination assay

Reactions were conducted with 100 nM of a fluorescein labeled 188 nt DNA substrate (Tri-Link Biotechnologies) with two 5’TTC motifs at 37°C in RT buffer (50 mM Tris, pH 7.5, 40 mM KCl, 10 mM MgCl_2_ and 1 mM dithiothreitol (DTT)). Reactions used 1000 nM A3C and were incubated for 0 to 60 min. The ssDNA substrate was previously reported (Ara et al., 2014). The reaction was initiated by the addition of ssDNA. Deamination reactions were stopped using phenol:chloroform extraction followed by two additional chloroform extractions. The deaminations were detected by treating the substrates with Uracil DNA Glycosylase (New England Biolabs) and heating under alkaline conditions. The ssDNA fragments were resolved on 10% (v/v) denaturing polyacrylamide gel. Gel photos were obtained using a Chemidoc-MP imaging system (Bio-Rad) and integrated gel band intensities were analyzed using ImageQuant (GE Healthcare).

Processivity reactions were carried out under single-hit conditions (<15% substrate usage) to ensure a single enzyme-substrate encounter (Creighton, Bloom, & Goodman, 1995). A processivity factor can be calculated under these conditions by comparing the quantified total amount of deaminations occurring at the two sites on the same ssDNA with a calculated theoretical value of deaminations assuming they were different deamination events (Chelico, Pham, Calabrese, & Goodman, 2006; Pham, Chelico, & Goodman, 2007). To determine these conditions, we determined the deamination over time and chose an appropriate time point. A processivity factor greater than 1.0 means the majority of double deaminations are catalyzed by a single enzyme, and therefore x-fold more likely to deaminate processively. A non-processive enzyme has a processivity factor of 1.0 or more commonly, does not have a visible amount of deamination at two sites under the single-hit conditions of the reaction. The specific activity was calculated under single-hit conditions by determining the picomoles of substrate used per minute for a microgram of enzyme.

### Steady-state rotational anisotropy

Steady-state rotational anisotropy reactions (60 μL) were conducted in buffer containing 50 mM Tris, pH 7.5, 40 mM KCl, 10 mM MgCl_2_, and 1 mM DTT and contained 10 nM of the fluorescein-labeled 118 nt ssDNA used for deamination assays. Increasing amounts of A3C was added (0 to 4500 nM). A QuantaMaster QM-4 spectrofluorometer (Photon Technology International) with a dual emission channel was used to collect data and calculate anisotropy. Measurements were performed at 21°C. Samples were excited with vertically polarized light at 495 nm (8-nm band pass), and vertical and horizontal emissions were measured at 520 nm (8-nm band pass). The apparent Kd was obtained by fitting to a single rectangular hyperbola equation using SigmaPlot version 11.2 software.

### Classical Molecular Dynamics

The initial structure of the human hA3C dimer was taken from the Protein Data Bank (PDB accession: 3VOW) and the peptide sequence was confirmed using Uniprot. The rhesus model (rhA3C) and all human-background variants were generated from the human dimer using Chimera to modify the peptide sequence. Four single mutants (R44Q, R45H, N115M, A144S), a double mutant (R44Q/R45H), and two triple mutants (R44Q/R45H/N115M, R44Q/R45H/A144S) were all generated from the human wildtype (WT) background. MolProbity and H++ were used for all systems to determine protonation states of amino acids at pH of 7.0 (Anandakrishnan, Aguilar, & Onufriev, 2012; Chen et al., 2010; Davis et al., 2007; Davis, Murray, Richardson, & Richardson, 2004; Gordon et al., 2005; Myers, Grothaus, Narayanan, & Onufriev, 2006). All models were neutralized to zero net charge with Cl^-^ and K^+^ counterions and solvated using TIP3P water with a 12Å minimum distance from protein surface to the edge of the solvent box (Jorgensen, Chandrasekhar, Madura, Impey, & Klein, 1983). This resulted in a solvated unit cell measuring 82Å x 84Å x 108Å. The FF14SB forcefield was used for the protein and TIP3P for water, Zn^2+^, and counterions (Jorgensen et al., 1983; Maier et al., 2015).

Molecular dynamics simulations were performed using OpenMM on XSEDE’s Comet HPC cluster (Eastman et al., 2017; Towns et al., 2014). Each system was equilibrated with iteratively reduced restraints on the protein to ensure stable simulation environment. The restraints began at 1000 kcal/mol and were reduced by half after every completed stage of equilibration until the restraint fell to below 1 kcal/mol. Each completed stage was run for a minimum of 1.0 ns (10^6^ timesteps) and checked for convergence to ensure stability of temperature (300K), density (1.0 g/mL), periodic box volume (approx 600 Å^2^), and total (potential and kinetic) energies of the system, resulting in a total equilibration time of 11.0 ns. To maintain active site geometry, harmonic restraints of 20 kcal/mol Å^2^ were applied between the active site zinc and E54 of each monomer subunit with an equilibrium distance of 5.0 Å. These restraints were maintained throughout the simulations. After equilibration, production dynamics were run for 100 ns using a 1 fs timestep and coordinates saved every 10 ps. A Langevin integrator was used as the thermostat with a Monte Carlo barostat in an NPT ensemble. The nonbonded cutoff distance was set to 10 Å, and the Ewald error tolerance was set to 10^-3^. All models were simulated in triplicate for a total of 300 ns of production.

Analysis of MD trajectories was performed using cpptraj (correlated motion, normal mode analysis, root mean squared deviation (RMSD) and fluctuation (RMSF), hydrogen bond interaction analysis) (Case et al., 2018). Data plots were generated with python, and 3D structure visualization was done using Chimera (Pettersen et al., 2004). The first 100 normal modes were calculated and their relative contribution to the total motion was examined. Dimer interaction energies were calculated using the AMBER-EDA program (available at https://github.com/CisnerosResearch/AMBER-EDA) by calculating the ensemble average Coulomb and Van der Waals interactions between each amino acid on one monomer with each amino acid on the other.

### Direct coupling analysis (DCA)

DCA is a global statistical inference model that is used to study coevolution in protein sequences (Morcos et al., 2011). Through DCA, an approximation of the global probability distribution for a multiple-sequence alignment creating a large quantity of homologous sequences can be modeled for a set of residual positions in the sequence. The DCA model can accurately estimate the direct covariations between any two variables while excluding secondary correlations between dependent variables. Therefore, DCA can be used to study molecular connectivity in various biological situations, such as the functionality and specificity for interacting proteins. DCA can also be used on non-sequence data, such as pharmacogenomic data (Jiang, Martinez-Ledesma, & Morcos, 2017).

In the DCA model, naturally occurring and modified protein sequences are assumed to be sampled from a Boltzmann distribution. Multiple sequences are grouped in a manner such that the homologous positions are aligned, forming a large sample space. The Boltzmann distribution is the most general and least-constrained model derived from maximum entropy modeling. DCA infers in an efficient manner the parameters of a large joint-probability distribution and uses these inferred parameters to determine estimates of the coupling between pairs of variables in such distribution. A metric to quantify such coupling between two positions in the alignment is called Direct Information (DI) (Morcos et al., 2011) which is zero if the two positions are uncoupled and positive otherwise. Higher DI values indicate a stronger dependency or functional relevance is present for the two residue sites. In this study, we utilize a metric called Cumulative Direct Information CDI*_i_*, which is the sum of the DI values for all the residue-residue pairs that a given residue *i* participates in a list of top *x* ranked pairs.

### Sequence Datasets

An NCBI multiple sequence alignment (MSA) was downloaded from the Pfam database for the APOBEC3 (PF18771) family (El-Gebali et al., 2019). Sequences with more than 27 (20% of the aligned sequence length) consecutive mispairing positions were removed. The resulting MSA was composed of 2504 sequences (Supplementary file 1). The MSA was processed using mean field DCA (mfDCA). The resulting DI pairs were then filtered for monomeric interactions and ranked from greatest to least DI value. A domain matching script using the FASTA sequence for hA3C (Q9NRW3) was used to match the calculated DI pairs to the original hA3C sequence.

### Molecular Simulation of Homodimeric Complexes driven by Evolutionary Couplings

The protein data bank (PDB) for hA3C (3VOW) was accessed to obtain all-atom coordinates of the protein crystal structure of hA3C (Kitamura et al., 2012). The structure was used to compute the solvent accessible surface area (SASA) of individual residues using *getarea* program (Fraczkiewicz & Braun, 1998). From the list of top coevolving DCA pairs, only those whose SASA summations were greater than 100 were taken for modeling studies to prioritize interactions at the protein surface. Using the PDB structure, all DCA pairs that were monomeric interactions, i.e., that contribute to the monomeric fold, were filtered out. From the remaining ranked DCA pairs, the top 25 were selected as dimeric interactions in a coarse-grain molecular simulation of the two interacting monomers.

The top 25 DCA pairs were incorporated as Gaussian potentials to drive formation of the hA3C dimer complex in a Structure Based Model (SBM)-MD simulation (dos Santos et al., 2015). The simulation process iteratively reduced the equilibrium distance between the monomer units to allow exploration of the dimer interface. A PDB file containing the two monomer subunits at 50Å was used as input in the SMOG server to generate parameter and topology files (Clementi, Nymeyer, & Onuchic, 2000; Noel, Whitford, Sanbonmatsu, & Onuchic, 2010). SBM potentials generated from the top 25 DCA pairs were inserted into the SMOG files. A version of GROMACS with support for SBM Gaussian potentials was used to run the molecular dynamics simulation (Lammert, Schug, & Onuchic, 2009). The final coordinates of the model were then compared to the initial PDB dimer structure.

### Contact Maps

The x-ray coordinates for hA3C (3VOW) were used to identify residue-residue physical contacts. A monomeric contact map was then created based on the proximity between amino acids of the same subunit, with a distance cutoff at 8Å between α carbon atoms. A dimeric contact map was generated based on the proximity between residues of different subunit, with a distance cutoff at 10Å between α carbon atoms. The two contact maps were then overlayed on top of each other to show all the residue-residue interactions in the PDB structure as well as the top 300 DI pairs calculated using DCA to give the final map.

### Ancestral reconstruction

Sequences of 18 primates were obtained from NCBI and were also described in (Wittkopp et al., 2016). Sequence of A3C from a New World Monkey, the white-faced capuchin, was used to root the tree of Old World Monkeys and Great Apes. Sequences were trimmed down to the coding region of A3C, gaps were removed, and the sequences at the ancestral nodes was calculated using FASTML (Moshe & Pupko, 2018). A translation alignment and a PHYML tree of 18 nucleotide sequences were generated in Geneious. The parameters for the FASTML run were as follows: model JTT; gammaon optimize; alignment status aligned; is jointon; branch optimization ontree build method RaxML.

## Acknowledgements

We thank Ossama Ibrahim for assistance with site directed mutagenesis. Research described in this paper was performed in part at the Protein Characterization and Crystallization Facility (PCCF), which is supported by the College of Medicine, University of Saskatchewan, Saskatoon, Canada. Reagents obtained through the NIH HIV Reagent Program, Division of AIDS, NIAID, NIH and Centre for AIDS Reagents, NIBSC, UK, were supported by EURIPRED (EC FP7 INFRASTRUCTURES-2012 - INFRA-2012-1.1.5.: Grant Number 31266). This work was supported by a Canadian Institutes of Health Research (CIHR) Grant PJT-162407 (L.C.), NIH R01GM108583 and NSF CHE-1856162 (G.A.C.), support to B.B. via NSF CHE-1757946, computing time from XSEDE TG-CHE160044 (G.A.C.), CASCaM with partial support from NSF CHE-1531468 (G.A.C.), NIH R35GM133631 and NSF MCB-1943442 (F.M.), NIH R01AI030927 (M.E.), D.W. was in the UW Post-Baccalaureate Research Education Program (PREP), and A.G. was supported by a Saskatchewan Health Research Foundation (SHRF) postdoctoral fellowship.

## Competing interests

All authors have no competing interests to declare.

## Notes

### Competing Interest Statement

The authors have declared no competing interest.

