## Supplementary Figures for "Divergence in dimerization and activity of primate APOBEC3C"

[1] Department of Biochemistry, Microbiology, and Immunology, College of Medicine, University of Saskatchewan, Saskatoon, Canada.
[2]Department of Chemistry, University of North Texas, Denton, Texas, USA.
[3]Department of Biological Sciences, Center for Systems Biology, University of Texas at Dallas, Richardson, Texas, USA.
[4]Division of Human Biology, Fred Hutchinson Cancer Research Center, Seattle, WA, U.S.A.
[5]Current Address: Department of Biochemistry & Biophysics, School of Medicine, University of North Carolina, Chapel Hill, North Carolina, U.S.A.
[6]Division of Basic Sciences, Fred Hutchinson Cancer Research Center, Seattle, WA, U.S.A.
[7]Department of Bioengineering, University of Texas at Dallas, Dallas, Texas, U.S.A.

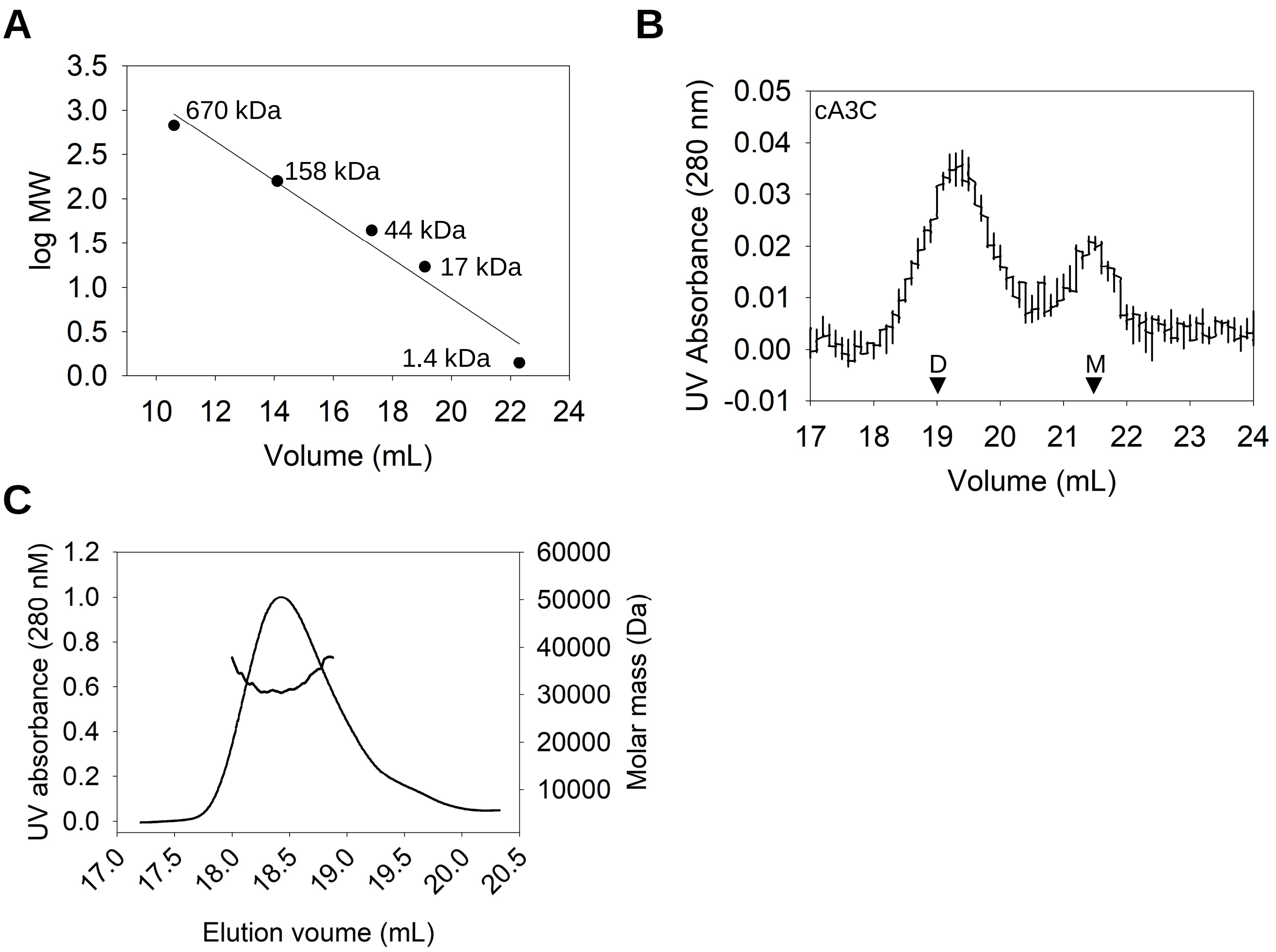


**Figure S1**. **Standards used for size exclusion chromatography. (A)** The standard curve generated from the G200 Increase column. Sizes correspond to Thyroglobulin (670 kDa), Gamma globulin (158 kDa), Ovalbumin (44 kDa), Myoglobin (17 kDa), and Vitamin B12 (1.4 kDa). **(B)** The cA3C is a mix of dimer (D) and monomer (M). The cA3C SEC was carried out as for rhA3C and showed that monomers elute at a volume greater than 20 mL and dimers elute at a volume smaller than 20 mL. **(C)** To confirm our measurements by SEC, which is based on standards, we also conducted multiangle light scattering (MALS) on the rhA3C R44Q/R45H/N115K, which had a single peak. We empirically calculated a molar mass of 30 kDa at the peak of elution, which is near the predicted molar mass from the amino acid sequence of 23 kDa.

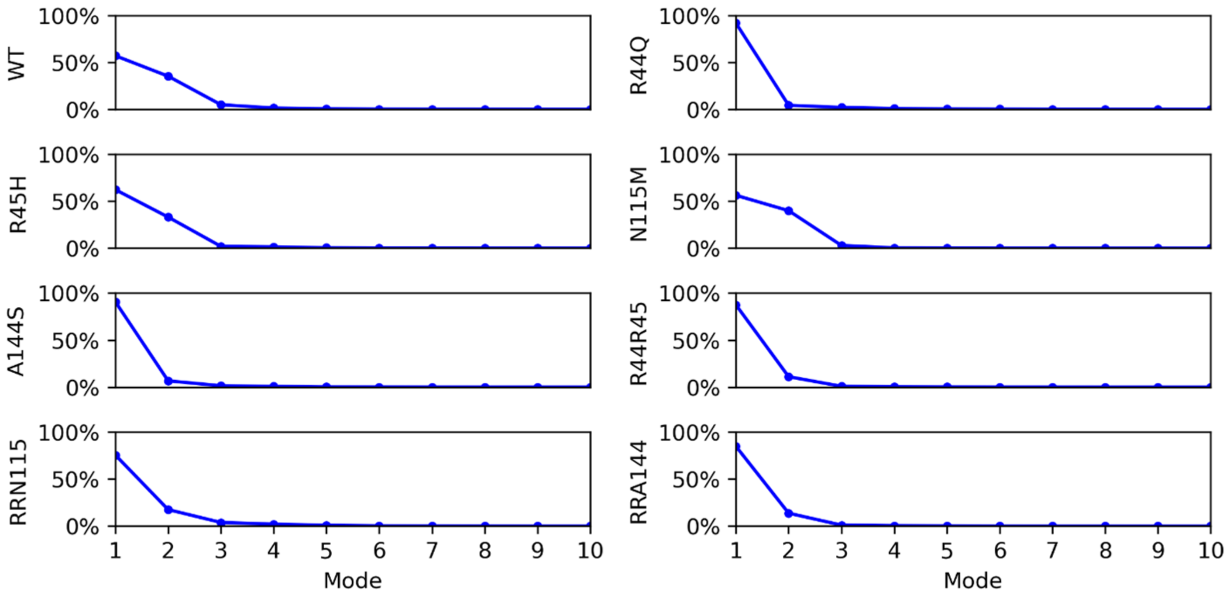


**Figure S2. Contribution to total dynamic motion from first ten (10) normal modes.** Amino acid changes are listed on the y-axis. R44R45 is the combined mutant R44Q/R45H. RRN115 is the combined mutant R45H/R44Q/M115N. RRA144 is the combined mutant R45H/R44Q/S144A.

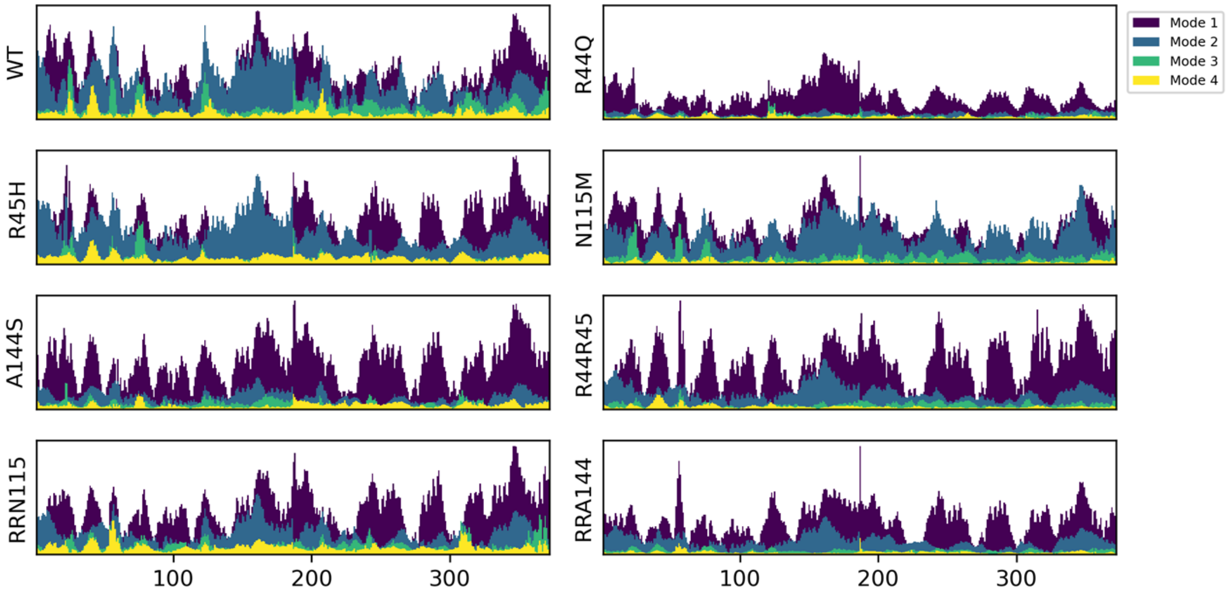


**Figure S3. First four (4) normal modes by residue.** As shown above, most of the essential motion is captured by the first three modes. The fourth mode is included here to show that it is negligible. R44R45 is the combined mutant R44Q/R45H. RRN115 is the combined mutant R45H/R44Q/M115N. RRA144 is the combined mutant R45H/R44Q/S144A.

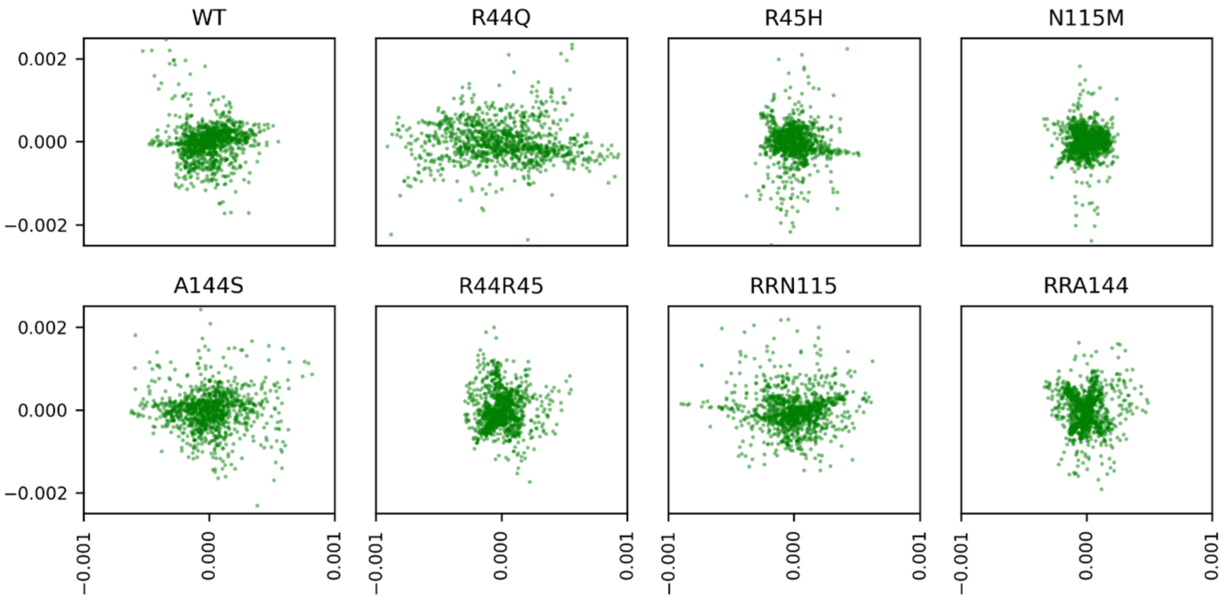


**Figure S4. PCA for first two normal modes.** R44R45 is the combined mutant R44Q/R45H. RRN115 is the combined mutant R45H/R44Q/M115N. RRA144 is the combined mutant R45H/R44Q/S144A.

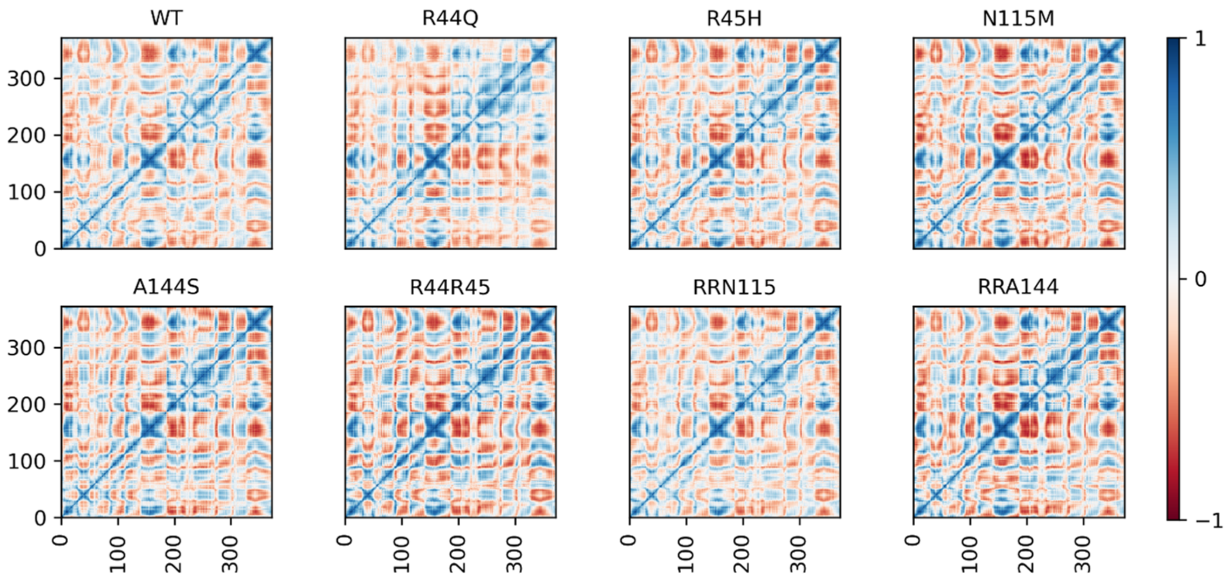


**Figure S5. Correlated motion matrices pairwise by amino acid.** R44R45 is the combined mutant R44Q/R45H. RRN115 is the combined mutant R45H/R44Q/M115N. RRA144 is the combined mutant R45H/R44Q/S144A.

**Table S1. Dimer Interaction Energy Decomposition Analysis.** Values shown as comparisons to wild type interaction energy.

| Mutations | ΔE compared to wild type (kcal/mol) |
|---|---:|
| R44Q | +161.6 |
| R45H | +384.4 |
| N115M | +173.8 |
| A144S | -13.0 |
| R44Q/R45H | +643.3 |
| R44Q/R45H/N115M | +631.3 |
| R44Q/R45H/A144S | +769.8 |

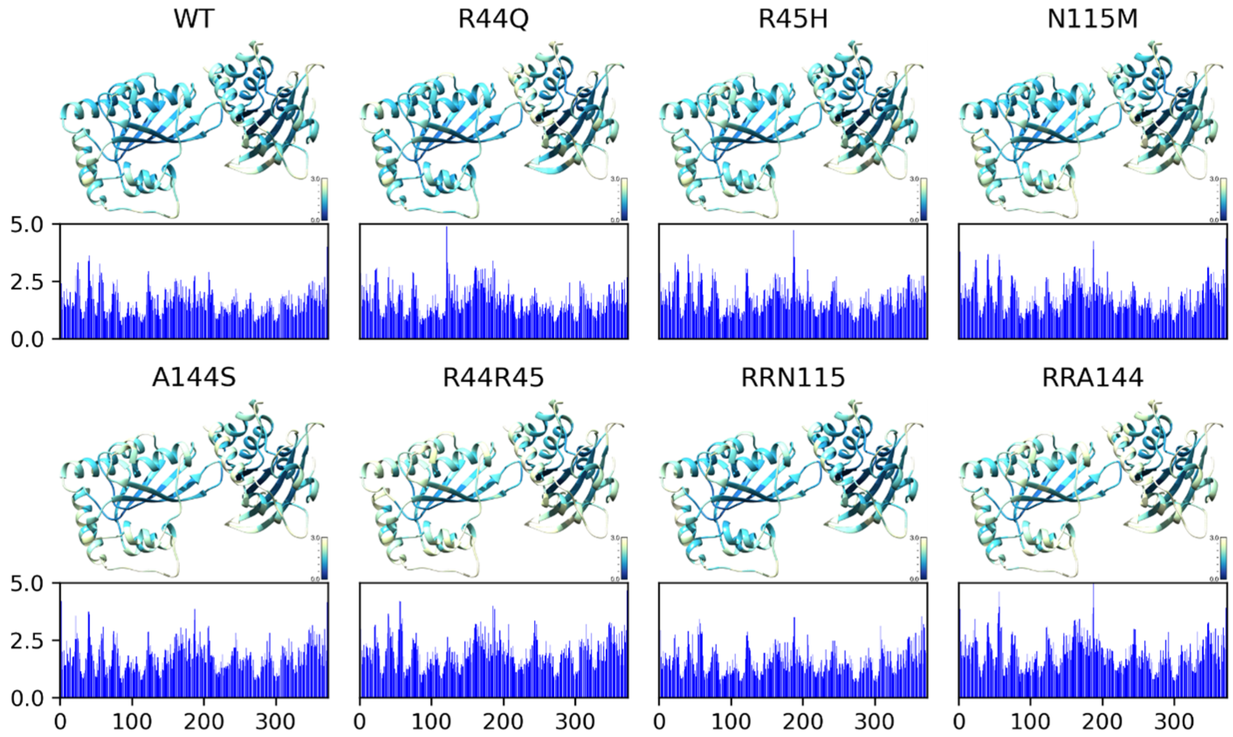


**Figure S6. RMSF by residue mapped to 3D structure.** Barplots below structures show raw RMSF data. R44R45 is the combined mutant R44Q/R45H. RRN115 is the combined mutant R45H/R44Q/M115N. RRA144 is the combined mutant R45H/R44Q/S144A.

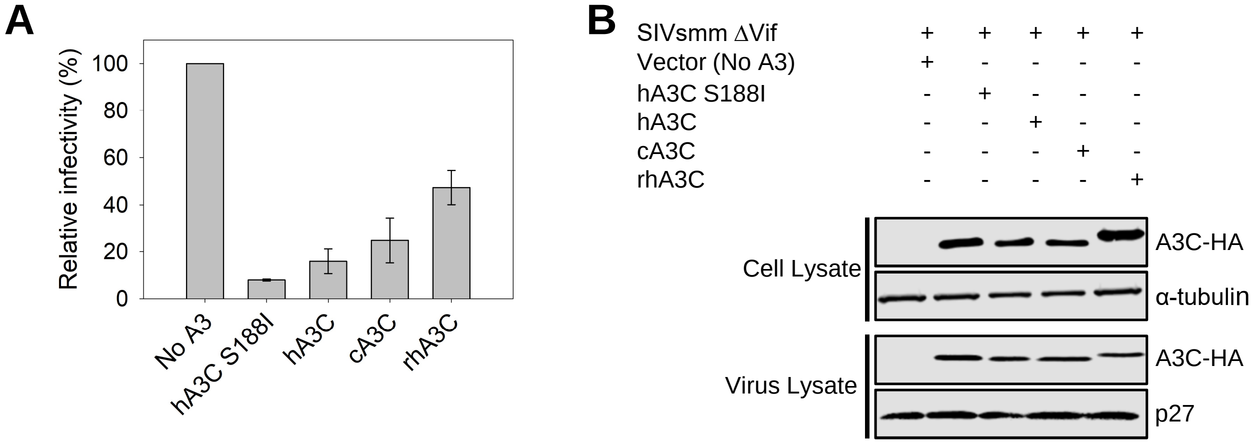


**Figure S7. A3C-mediated restriction of SIVsmm. (A)** SIVsmm L5 ΔVif infectivity was measured by β-galactosidase expression driven by the HIV-1 5´LTR from TZM-bl cells infected with SIVsmm that was produced in the absence or presence of 3xHA tagged hA3C, hA3C S188I, cA3C, and rhA3C. Results normalized to the no A3 condition are shown with the Standard Deviation of the mean calculated from at least three independent experiments. (**B**) Immunoblotting for the HA tag was used to detect A3C enzymes expressed in cells and encapsidated into SIVsmm ΔVif virions. The cell lysate and virion loading controls were α-tubulin and p27, respectively.

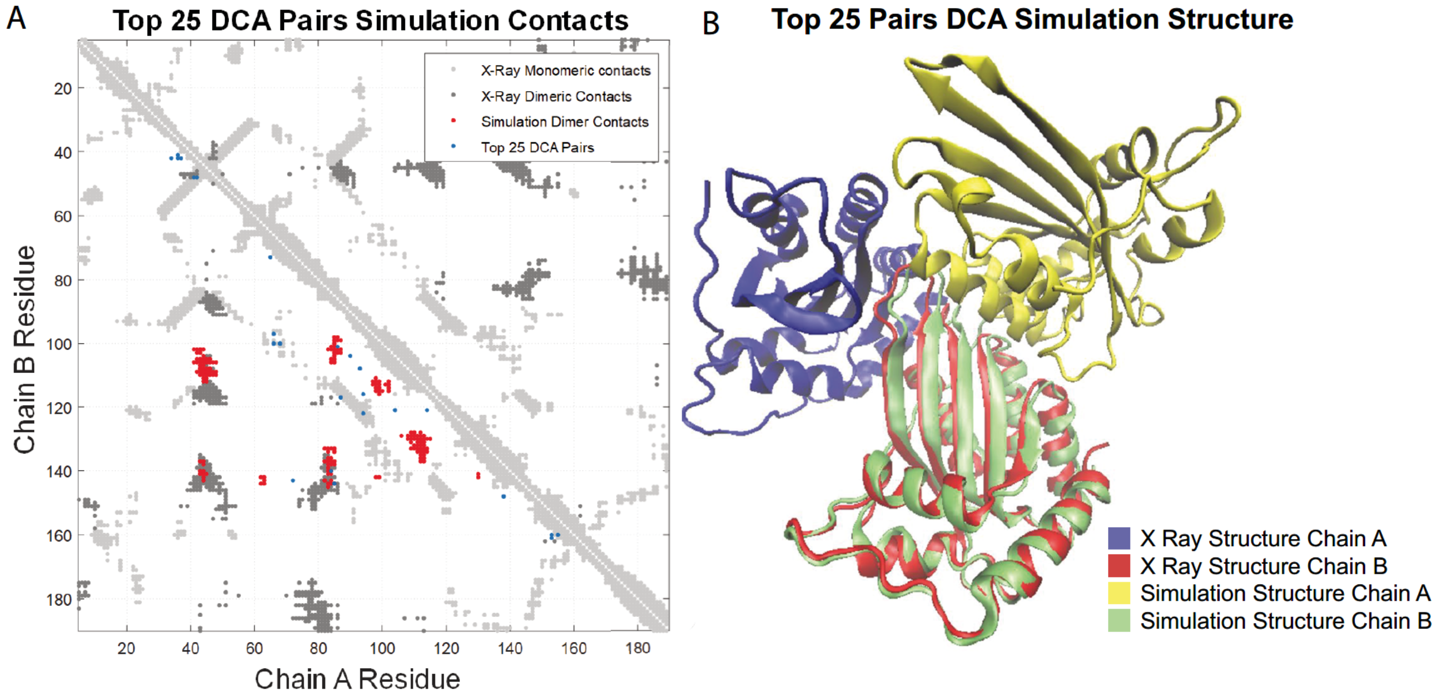


**Figure S8. Structure Based Model molecular dynamics simulation driven by coevolved pairs is used to predict dimeric complexes.** A simulation utilizing the top dimeric residue-residue couplings estimates a putative dimeric complex. (A) shows a comparison of the dimeric interactions found in the 3VOW crystal structure (dark grey dots) and the dimeric interactions identified by the homodimer prediction (red dots). Although the predicted complex and the crystal structure (B) are not identical, they do share several important dimeric interactions, particularly those involving the residue 144.

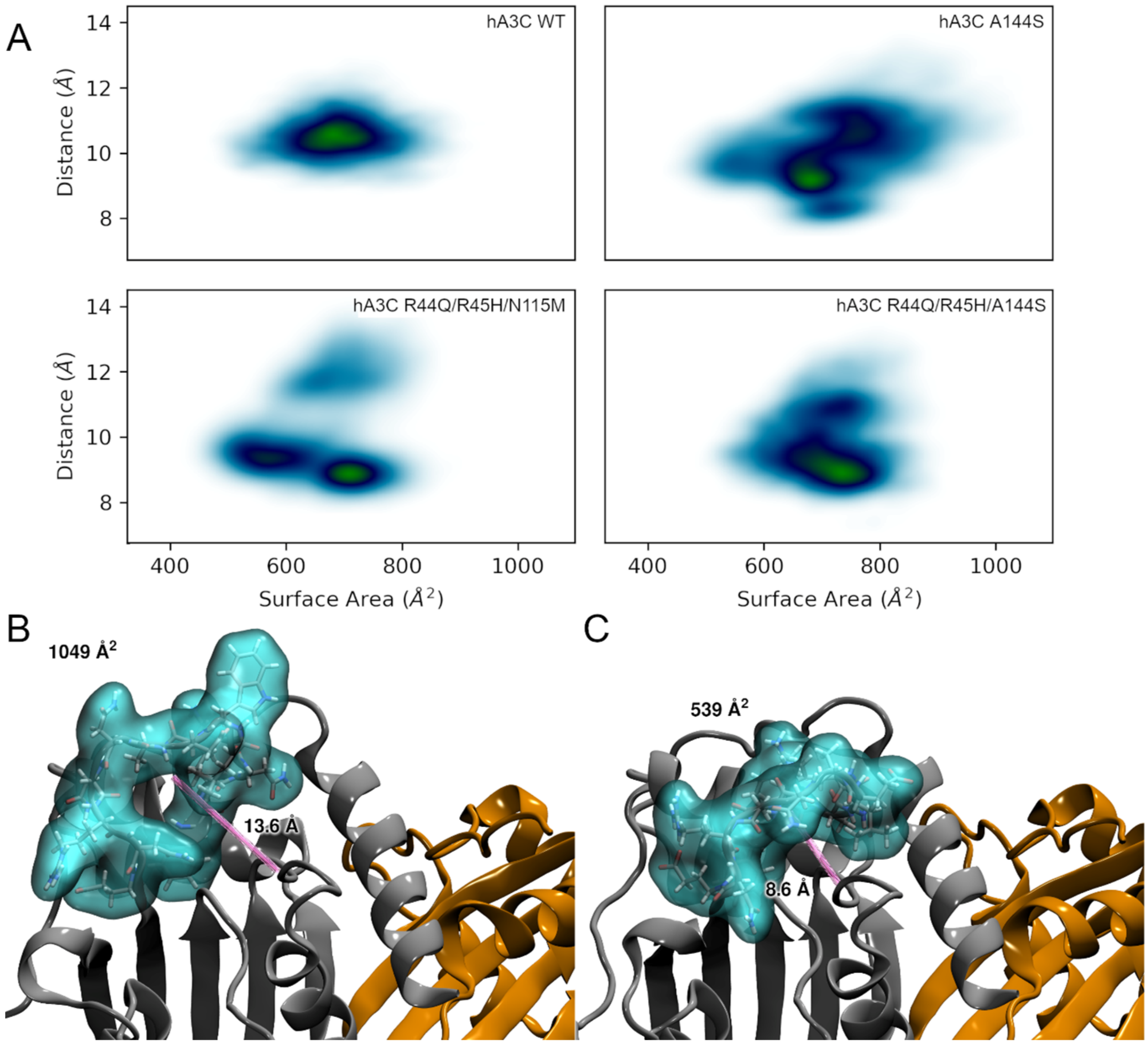


**Figure S9. Analysis of loop 1. (A)** Density of states for hA3C WT, A144S, R44Q/R45H/N115M, and R44Q/R45H/A144S over time comparing solvent accessible surface area (SASA) of loop 1 residues (positions 21-32) and distance from center of mass of α-carbons of loop 1 and center of mass of α-carbons of residues on helix 4 (positions 121 to 128), positioned across the active site from loop 1. These measurements were used to determine the approximate openness of the active site. The upper right quadrant in the plots indicates states with a more open active site, and points in the lower left quadrant indicates states with a more closed active site. Vertical spread indicates the range of distance over the simulation time, and horizontal spread indicates the amount of loop 1 that is extended into solvent, another measure of openness. The hA3C WT shows a higher density region around the center of the plot. The hA3C A144S exhibits considerable differences compared with the hA3C WT, including separated regions of density which suggest distinct "locked" states in which the systems spend proportionally more time. **(B)** hA3C WT with loop 1 (blue surface) in an open conformation with side chains extending away from the protein surface and distance (pink line) across active site, **(B)** A3C with loop 1 (blue surface) in closed conformation with side chains closer to the protein surface and distance (pink line) across active site.

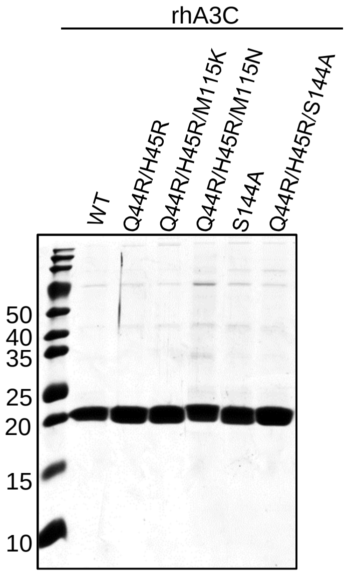


**Figure S10. Purified rhA3C.** Five micrograms of rhA3C WT and mutants were resolved by SDS-PAGE.
